# Dynamic rewiring of electrophysiological brain networks during learning

**DOI:** 10.1101/2022.04.08.487166

**Authors:** Paolo Ruggeri, Jenifer Miehlbradt, Aya Kabbara, Mahmoud Hassan

## Abstract

Human learning is an active and complex process. However, the brain mechanisms underlying human skill learning and the effect of learning on the communication between brain regions, at different frequency bands, are still largely unknown. Here, we tracked changes in large-scale electrophysiological networks over a 6-week training period during which participants practiced a series of motor sequences during 30 home training sessions. Our findings showed that brain networks become more flexible with learning in all the frequency bands from theta to gamma ranges. We found consistent increase of flexibility in the prefrontal and limbic areas in the theta and alpha band, and over somatomotor and visual areas in the alpha band. Specific to the beta rhythm, we revealed that higher flexibility of prefrontal regions during the early stage of learning strongly correlated with better performance measured during home training sessions. Our findings provide novel evidence that prolonged motor skill practice results in higher, frequency-specific, temporal variability in brain network structure.

**AUTHOR SUMMARY:** We investigated the large-scale organization of electrophysiological brain networks of a cohort of 30 participants practicing a series of motor sequences during 6 weeks of training. With learning, we observed a progressive modulation of the dynamics of prefrontal and limbic regions from theta to alpha frequencies, and of centro-parietal and occipital regions within visuomotor networks in the alpha band. In addition, higher prefrontal regional flexibility during early practice correlated with learning occurring during the 6 weeks of training. This provides novel evidence of a frequency-specific reorganization of brain networks with prolonged motor skill learning and an important neural basis for non-invasive research into the role of cortical functional interactions in (visuo)motor learning.

## 1. INTRODUCTION

Learning is a multidimensional concept with various definitions depending on the realm and the level of investigation. At the most fundamental level, learning results in brain architecture modifications through modulation of neural synapses (Smolen et al., 2016) causing changes in neural information processing (Dayan and Cohen, 2011). Behaviorally, such adaptability allows transforming an initially cognitively demanding and slow activity into a more spontaneous and automatic process (Grafton et al., 2008). The last two decades have been characterized by a growing interest in the scientific community in the area of learning, particularly motor skill learning. Several previous learning experiments with MRI produced in the context of pre/post training regimes were able to clearly demonstrate that macro-scale investigations through brain structural and functional connectivity analyses can successfully identify learning-related changes in cortical and subcortical brain networks. More specifically, structural investigations (e.g., Draganski et al., 2006; Sampaio-Baptista et al., 2014; Taubert et al., 2011, 2010) have consistently reported associations between gray matter increase and learning in task relevant areas including motor, parietal, and prefrontal cortex. On the other hand, following the practice of various motor learning tasks, functional connectivity (e.g., Mehrkanoon et al., 2016; Sami et al., 2014; Sampaio-Baptista et al., 2015; Sun et al., 2006; Taubert et al., 2011; Tung et al., 2013) was seen to be modulated within numerous cortical (motor, frontoparietal, sensorimotor, and visual) and sub-cortical (within and between cerebellar nuclei, thalamus and basal ganglia) networks as well as between cortical and sub-cortical areas.

However, the entire field still remains underrepresented, probably because exploring learning-related aspects requires long-term longitudinal designs (from weeks to months of practice) and consequently considerable deployment of research material and human know-how. In addition, there is a need for studies dedicated to the temporal dynamics of interactions between brain regions during learning phases that can surpass and augment knowledge provided by - mostly resting-state - pre/post investigation protocols. In this regard, the most significant advances have been recently achieved using fMRI techniques and the joint development and use of innovative data analysis methodologies (Mucha et al., 2010) perfectly tailored to characterize brain dynamic processes such as those underlying learning. Surprisingly, studies using electrophysiological neuroimaging techniques such as magneto/electro-encephalography (M/EEG) are rare and most of the few that do exist have investigated motor adaptation (e.g., Gentili et al., 2015; Mehrkanoon et al., 2016; Miraglia et al., 2018) and, to a lesser extent, motor skill learning (e.g., Schubert et al., 2021; Tzvi et al., 2018, 2016) during very short periods of practice (from single sessions to very few days). This is despite the flexibility and ease of use of these neuroimaging techniques and the recent development and adaptation of existing brain dynamics methodologies to M/EEG data (Hassan and Wendling, 2018; O’Neill et al., 2017; Tabbal et al., 2021).

The combination of fMRI techniques and the development of powerful mathematical tools from network science and graph theory (Barabási, 2013; Stam, 2014) to identify metrics assessing brain network dynamics have allowed for the characterization of changes in region-to-region interactions in the brain (Bassett and Sporns, 2017; Kivelä et al., 2014; Mucha et al., 2010). This approach provided a breakthrough in the description of adaptive fundamental brain processes supporting learning (Bassett et al., 2011, 2013b, 2015; Mattar et al., 2018; Reddy et al., 2018). Most of the investigations were based on the use of learning tasks involving motor and visual systems and the acquisition of new motor skills in particular. Results of these studies have shown that learning is accompanied by large-scale dynamic variations over timescales of seconds to minutes (Allen et al., 2014; Betzel et al., 2017; Zalesky et al., 2014) across distributed networks associated with executive functions, visual processing and motor execution during task performance (Bassett et al., 2011, 2013b, 2015). Compared to classical approaches focused on the activation strength of isolated brain regions, these studies brought specific insights into how broad-scale interconnected systems accommodate learning. In particular, they showed that the recruitment of core visual and motor areas did not vary with practice, while their integration decreased with training. In addition, they revealed that individual learning was related to the extent of the decrease in the integration across medio-frontal brain areas required in higher order cognitive processes (Bassett et al., 2015). These findings were accompanied by evidence that brain network community structures (Mattar and Bassett, 2016) (i.e., densely connected regions within a network) are not stable over time but evolve dynamically. In the context of motor skill learning, several studies have assessed brain network dynamics by quantifying the flexibility of brain regions in changing their community allegiance over time (Bassett et al., 2011, 2013b; Betzel et al., 2017; Reddy et al., 2018). From a computational point of view, flexibility is commonly used as a tool to determine the temporal variability of community structures and quantifies the frequency that a brain region changes its modular affiliation over time. High values of flexibility are interpreted as indicative of continuous change in community affiliation, while lower values are indicative of stable affiliation over time (Bassett et al., 2011, 2013b; Betzel et al., 2017). In addition, it has been demonstrated that regions with low (high) flexibility tend to be strongly (weakly) -connected network nodes (Bassett et al., 2013b). For motor skill learning, this approach was particularly suited to describe the transition from controlled towards automatic processes (Bassett et al., 2011, 2013b; Reddy et al., 2018). Bassett and colleagues (Bassett et al., 2011) were the first to highlight how brain flexibility was temporally correlated with subsequent performance during the acquisition of a simple motor skill, and to suggest a possible modulation of flexibility with learning. Later, using a longitudinal motor skill learning protocol with fMRI, the same authors (Bassett et al., 2013b; Reddy et al., 2018) confirmed the relationship between performance and flexibility and the increase of flexibility with learning. Brain network flexibility has also been shown to correlate with cognitive flexibility (Mattar and Bassett, 2016), positive affect, surprise and fatigue (Betzel et al., 2017). Taken together, this complementary evidence highlights the advantage of using brain network flexibility to characterize the complex dynamics of brain networks.

The EEG literature on motor skill learning lacks brain network studies exploring the time hierarchy and dynamics of learning-related brain functions over longer period of motor skill acquisition, but provides important research showing that the specific recruitment of brain regions during performance of visuomotor adaptation and motor skill learning tasks takes place in frequency bands from theta to gamma ranges. Several studies have shown EEG power modulations in areas relevant to the performance of these tasks over frontal, motor and visual areas. More specifically, the involvement of prefrontal areas, often implicated in aspects of executive control with activity modulated by attentional and cognitive load (Klimesch, 1999; Langer et al., 2013; Scheeringa et al., 2008), has been mainly observed in the theta (Aliakbaryhosseinabadi et al., 2021; Crivelli-Decker et al., 2018; Koch et al., 2020) frequency band and, to a lesser extent, in the alpha (Meyer et al., 2014), beta (Aliakbaryhosseinabadi et al., 2021; Jahani et al., 2020), and gamma (Aliakbaryhosseinabadi et al., 2021) bands. Others have shown the involvement of sensorimotor and motor areas in the beta (Aliakbaryhosseinabadi et al., 2021; Andres and Gerloff, 1999; Aoki et al., 2001; Boonstra et al., 2007; Cunha et al., 2006; Heinrichs-Graham et al., 2016; Jahani et al., 2020; Jerbi et al., 2004; Pollok et al., 2014; Rilk et al., 2011) and alpha (Andres and Gerloff, 1999; Jerbi et al., 2004; Meyer et al., 2014; Rilk et al., 2011; Rueda-Delgado et al., 2019; Schubert et al., 2021; Tzvi et al., 2018, 2016; Zhuang et al., 1997) bands, while the implication of the theta (Koch et al., 2020; Perfetti et al., 2011; Studer et al., 2010; Tzvi et al., 2016) and gamma (Aliakbaryhosseinabadi et al., 2021; Aoki et al., 2001; Perfetti et al., 2011) bands was less consistently reported. A number of studies have also highlighted the recruitment of visual processing areas during adaptation and visuomotor sequence tasks in the alpha (Meyer et al., 2014; Rilk et al., 2011; Tzvi et al., 2018, 2016) band and, less consistently, in the beta (Rilk et al., 2011) and gamma (Tzvi et al., 2016) bands. Important evidence is provided by connectivity studies during visuomotor practice. With respect to an executive network represented by functional interactions between prefrontal, central and parietal regions, Rilk and colleagues (Rilk et al., 2011) revealed an association between frontocentral coupling in the alpha band and increased errors in a tracking task. In line with this finding, a decrease in coherence between prefrontal and central regions was observed with ongoing learning and improved performance in the theta, alpha and beta bands (Gentili et al., 2015). Along the same lines, it was suggested that this network becomes less relevant with acquired encoding (Tzvi et al., 2018). With respect to a visuomotor network formed by functional interactions between central, parietal and occipital regions, it was observed that the coherence between motor and visual regions in the alpha and beta band increased during visuomotor task performance (Classen et al., 1998; Erla et al., 2012). Consistently, it was shown that coherence in these regions increased during task execution in the beta band and that it was related to better tracking task performance in the alpha band (Rilk et al., 2011). Finally, few brain network studies showed an increase of modularity and transitivity metrics with learning, in the theta and alpha bands (Miraglia et al., 2018), and a relationship between small worldness and better learning in the alpha band (Vecchio et al., 2018).

Taken together, this substantial evidence from neuroimaging studies highlights how there is still much to be clarified about the mechanisms underlying motor skill learning and how functional communication between different brain regions is modulated by learning. In particular, to provide a more detailed description of the dynamics of these functional brain networks, much work is needed to reveal how functional connectivity modulations at different frequencies occur over extended periods of practice. Here, we aimed to close this gap by leveraging on an EEG-based time varying connectivity analysis in the context of a longitudinal motor skill protocol. To this end, inspired by previous fMRI studies on this topic (e.g., Bassett et al., 2013), we investigated whether learning a new motor skill from visual cues was accompanied by a reorganization of brain networks also visible in the EEG. In particular, we assessed whether extended practice was accompanied by frequency specific modulations of flexibility across brain regions and whether the dynamics of brain networks in early stages of learning was related to motor skill performance (Bassett et al., 2011, 2013b; Mattar et al., 2018). We studied a cohort of healthy adult human subjects (*N* = 30) who performed thirty sessions of a discrete sequence-production task (DSP) over 6 weeks (Figure 1). During this period, participants underwent four High Density (HD)-EEG sessions while practicing the same DSP task, following an experimental protocol used in several previous works (Bassett et al., 2013b, 2015; Mattar et al., 2018; Reddy et al., 2018; Wymbs and Grafton, 2015). We examined network reconfiguration using dynamic network measures involving community detection and flexibility analysis (Bassett et al., 2013b, 2013a; Mucha et al., 2010).

**Figure 1:**
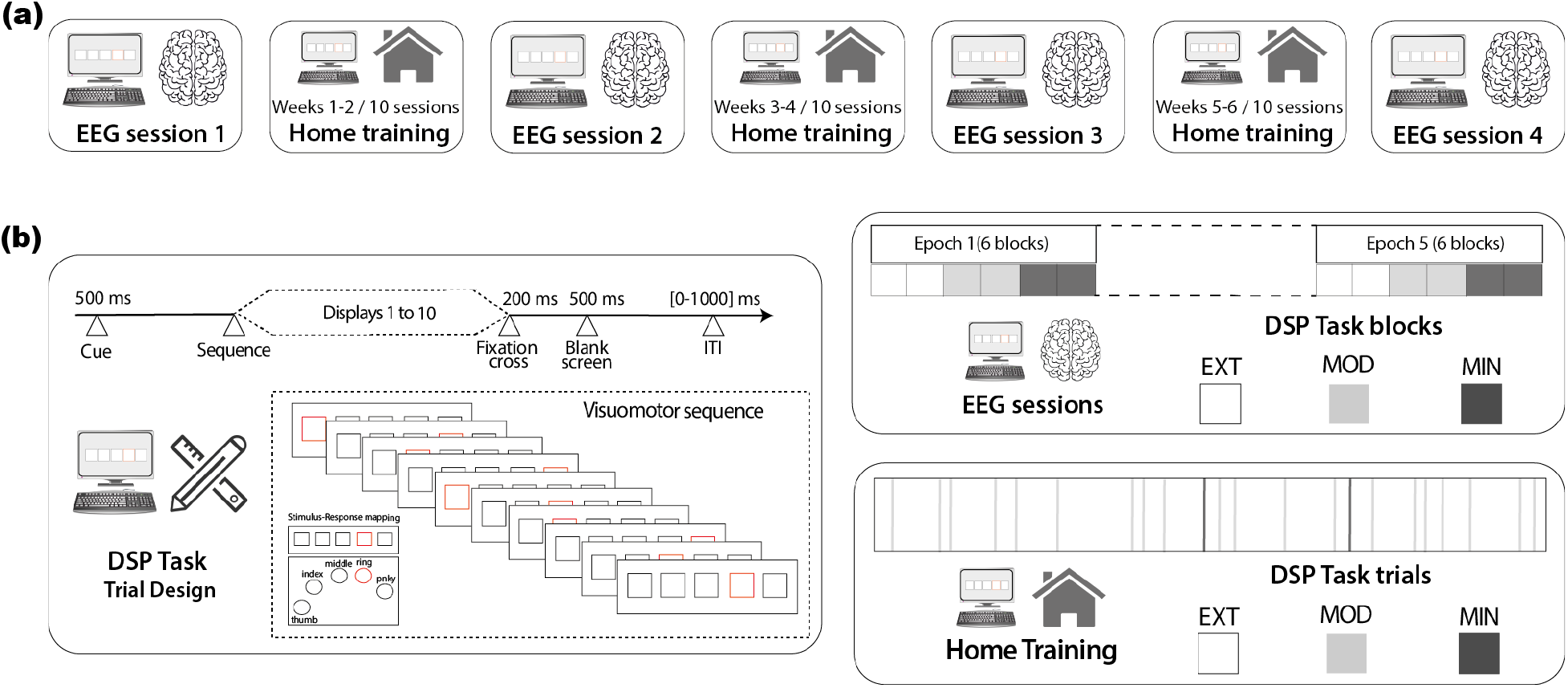
Experimental protocol and task design. (a) EEG sessions in the laboratory were interleaved with training sessions at home. Study participants practiced the DSP task for the first time during EEG session 1 and returned for another EEG session every 2 weeks (EEG sessions 2 to 4) after ten home training sessions of the DSP task; (b) On the left, the trial design of the DSP task. The numbers represent the duration of the presented stimuli consisting of a 500 ms cue preceding the practice of the visuomotor sequence made of 10 consecutive displays, followed by a fixation cross (200 ms), a blank screen (500ms) and an inter-trial interval of varying duration (0-1000ms). To terminate the sequence, participants had to select the correct stimulus-response mapping. On the right, the DSP task practiced during EEG and home training sessions. The former consisted of 5 epochs made of 6 blocks (2 for EXT, MOD and MIN). Within each block, 5 sequences of each type (EXT 1 & 2, MOD1 & 2, and MIN 1 & 2) were practiced in respective trials. The appearance of the sequences was randomized within each block and the appearance of the blocks was randomized within each epoch. At the end of each EEG session, participants practiced 100 EXT trials (50 EXT1 & EXT2), 100 MOD trials (50 MOD1 & MOD2), and 100 MIN trials (50 MIN1 & MIN2). The DSP task practiced during the home sessions consisted of 150 trials, divided into 64 trials for each EXT sequence, 10 trials for each MOD sequence, and 1 trial for each MIN sequence. Trials were randomized.

We hypothesized a (i) global increase in flexibility (Bassett et al., 2013b; Reddy et al., 2018) over successive EEG sessions. More specifically, we (ii) expected that a reduced involvement of prefrontal regions in cognitive control processes with learning (Bassett et al., 2015) would manifest in an increased availability of interaction with other regions and thus in the form of increased flexibility of regions of this network in theta (Aliakbaryhosseinabadi et al., 2021; Gentili et al., 2015; Miraglia et al., 2018), alpha (Gentili et al., 2015; Miraglia et al., 2018; Rilk et al., 2011) and, to a lesser extent, beta (Gentili et al., 2015) frequency bands. Furthermore, we hypothesized that (iii) the previously observed learning-related reduction in integration between regions in the somatomotor and visual networks (Bassett et al., 2015) may manifest in the form of increased flexibility of regions belonging to these networks predominantly in the alpha and beta frequency bands (Classen et al., 1998; Erla et al., 2012; Rilk et al., 2011). Lastly, based on evidence that both the functional disengagement of prefrontal regions (Bassett et al., 2015) and the configuration of brain networks during resting state (Mattar et al., 2018) and early phases of learning (Bassett et al., 2013b, 2011) are related to the amount of learning in subsequent practice sessions, (iv) we expected a positive relationship between early prefrontal regions flexibility and the learning rate over 6 weeks of practice in the theta, alpha and beta frequency bands. Finally, it is important to note that the methods used in this work, although extensively used and representing the methodological reference of the totality of the works in the field, are still undergoing continuous improvement. To shed light on possible confounding factors (such as the signal’s power and the task design) that could influence the extracted metrics, these methods have been put in perspective and further explored in a dedicated section in the supplementary material (section 4).

## 2 RESULTS

We investigated the dynamics of brain network reconfiguration underlying prolonged training of a simple motor skill (Figure 1 and 2). We focused on functional connectivity between different brain regions at four frequency bands (Theta: 4-8 Hz; Alpha: 8-12 Hz; Beta: 12-28 Hz; Gamma: 28-45 Hz) measured during 4 EEG sessions. These sessions were executed every 2 weeks as part of an experimental protocol quantifying learning of a simple sensorimotor sequence learning task practiced during 30 home-based sessions over 6 weeks (Figure 1a). During this period, 30 participants learned six 10-element sequences, practiced in pairs at an intensive (two EXT sequences: EXT1 & EXT2), moderate (two MOD sequences: MOD1 & MOD2) and minimal (two MIN sequences: MIN1 & MIN2) pace (Figure 1b). To characterize brain dynamics (Figure 2), functional connectivity matrices were computed from the EEG source-localized signals extracted during the execution of each trial of the task (corresponding to the execution of a sequence). Multilayer temporal networks (Holme and Saramäki, 2012) were then separately built for each training intensity (EXT, MOD and MIN) by concatenating, in successive layers, connectivity matrices corresponding to consecutive trials of the same type (EXT, MOD or MIN). These multilayer temporal networks were then analyzed using computational tools for dynamic community detection (Bassett et al., 2013a; Mucha et al., 2010) consisting in the maximization of a multilayer modularity quality function *Q* describing the within- and across-layers relationships between brain regions. This was done to identify functional modules (i.e., clusters of communities of brain regions sharing correlated brain dynamics) across the entire course of learning. Each community of brain regions can be thought of as segregated network coding for specific cognitive or motor functions. To quantify the dynamic properties of community structure and capture changes over the 6 weeks of continuous learning, we used flexibility (Bassett et al., 2013b, 2011) as network diagnostics. Finally, behavioral task performance was quantified by the time required to complete an entire sequence (movement time - MT). Following the approach validated in previous studies (Bassett et al., 2015, 2013b; Mattar et al., 2018), the reduction in MT with ongoing practice was used as an indicator of learning.

**Figure 2:**
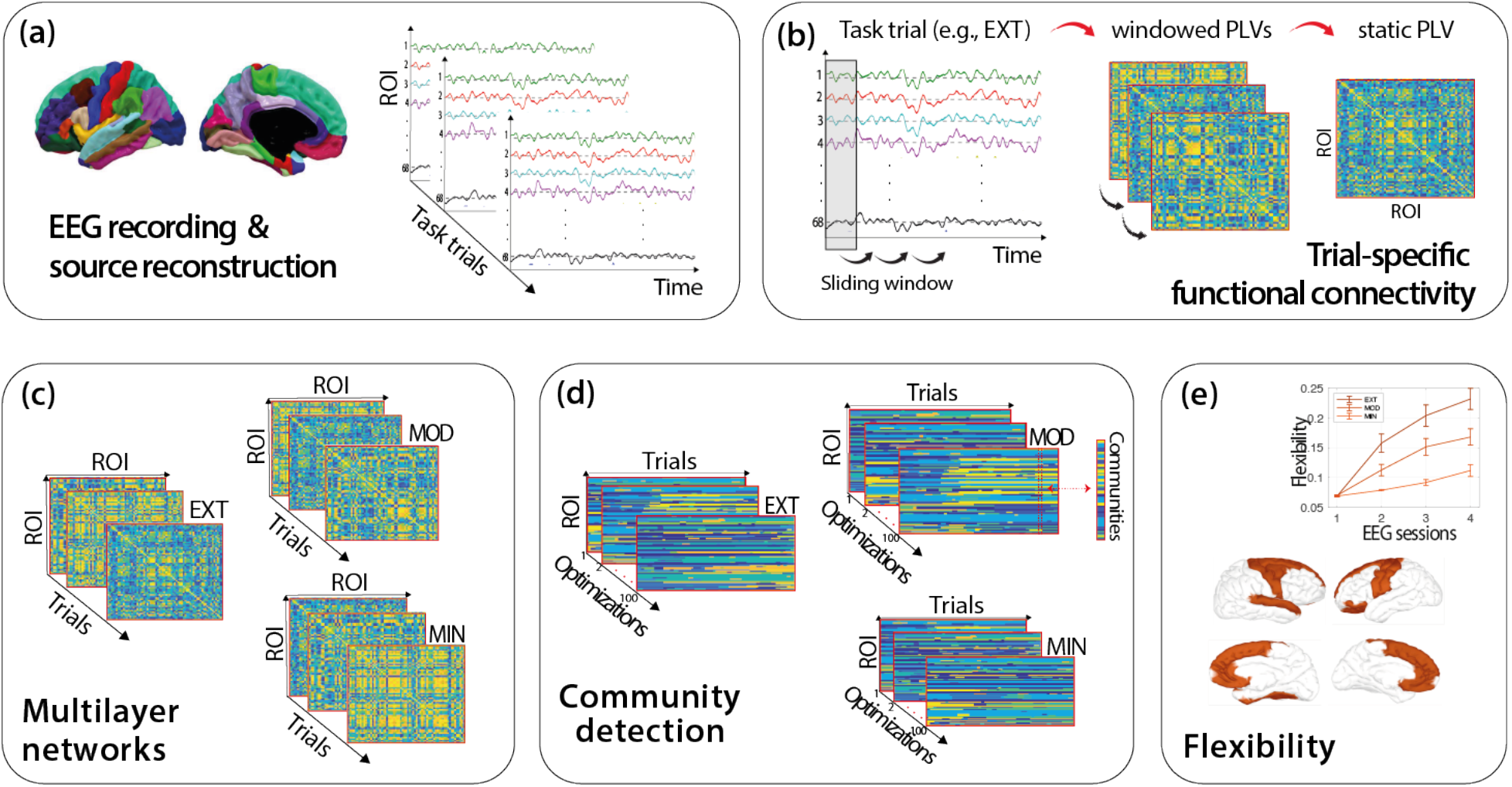
Methodological framework to compute flexibility. (a) EEG data were recorded and the source activity of 68 ROI of the Desikan-Killiany atlas in each trial of the DSP task was computed using the wMNE algorithm; (b) Trial-specific functional connectivity was obtained as a static PLV matrix obtained by averaging the windowed PLVs obtained with sliding window approach where connectivity was computed within each temporal window; (c) Multilayer network tensors for each type of sequence (EXT, MOD, and MIN) were built by concatenating static PLVs from consecutive trials of the same type; (d) Community detection was implemented on each multilayer network tensor through the modularity maximization method. Modularity maximization was run 100 times, leading to 100 optimized community assignments for each sequence type; (e) Global and regional flexibility was computed for each run and then averaged over the 100 runs. The procedure outlined above was applied to the EEG signal of each participant in each frequency band (theta to gamma) and experimental session (1 to 4).

### 2.1 Behavioral performance

We first sought to determine, from a behavioral point of view, the effectiveness of the experimental manipulation as practice progressed. To this end, changes in MT (i.e., time required to practice a sequence correctly) across the six weeks of training were assessed with repeated measures ANOVAs (with training intensity -EXT, MOD and MIN- and laboratory session - 1 to 4 - as within-subject factors; Figure 3a-b). MT significantly decreased across sessions (p < .001) and differed between EXT, MOD and MIN sequences (p < .001) with lowest values observed during the execution of the EXT sequences (EXT: M = 2.22 s, SE = 0.11 s; MOD: M = 2.54 s, SE = 0.13 s; MIN: M = 3.26 s, SE = 0.12 s). MT decreased more rapidly across scan sessions for sequences that were extensively practiced (EXT) as compared to less practiced ones (interaction effect, p < .001). Taken together, these analyses showed that the MT recorded during EEG sessions consistently decreased within-subjects for all practiced sequences (see also Figure 3b for a qualitative appreciation of this reduction as the subject-specific level), corroborating previous findings (Bassett et al., 2015; Wymbs and Grafton, 2015).

**Figure 3:**
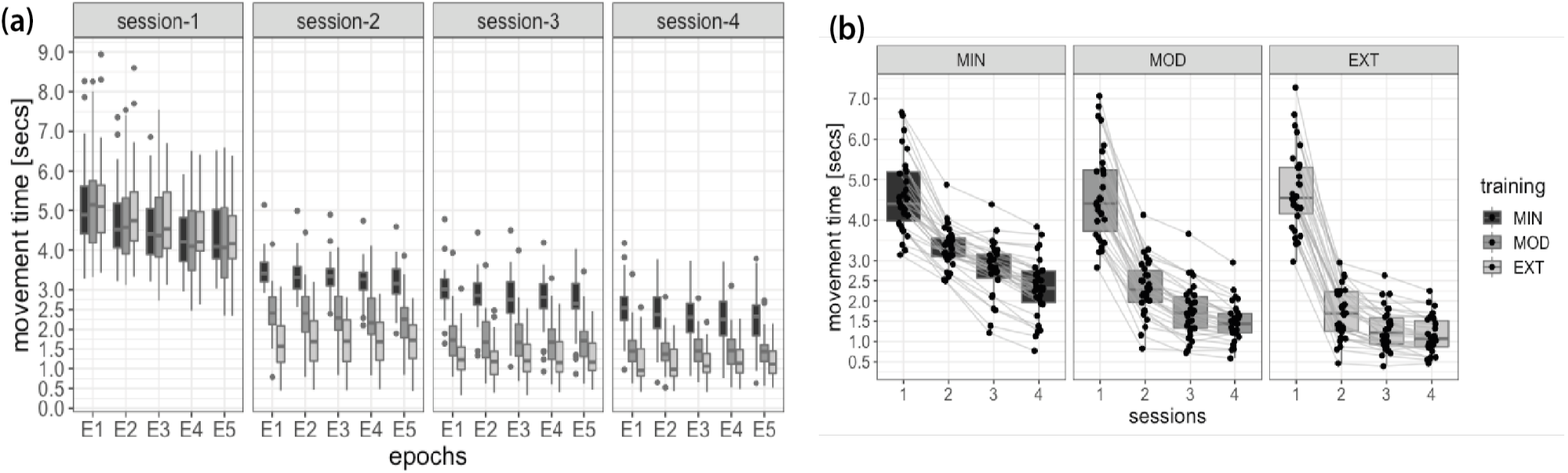
Behavioral indicators of performance. (a) MT recorded during EEG sessions and grouped by training intensity (EXT, MOD and MIN), revealing different improvements as a function of practice exposure during home sessions. During each EEG session the EXT, MOD and MIN sequences are equally practiced during 5 consecutive epochs. (b) Paired observations highlighting the distribution of MT for EXT, MOD and MIN sequences recorded during EEG sessions 1-4, and revealing that all participants decreased their MT during the 6 weeks of practice. Lower and upper box boundaries represent the 25th and 75th percentiles (Q1 and Q3); the difference between Q1 and Q3 is the interquartile range (IQR); the horizontal line a inside the box represents the median of the distribution; the lower and upper extreme lines show Q1-1.5xIQR and Q3+1.5xIQR, respectively; finally, the filled gray circles are data points falling outside the lower and upper extreme lines.

Following previous works using the same experimental procedure (Bassett et al., 2015, 2013b; Mattar et al., 2018; Reddy et al., 2018), we quantified learning by assessing the decrease in the MT required to correctly perform each sequence as practice progressed. The learning rate *K* of each participant was computed as the exponential drop-off of MT (*r*^*2*^ : M = 0.96; SD = 0.05) required to perform the two EXT sequences during the 30 home-sessions (see Methods section and Figure S1). Indeed, these sequences were extensively practiced (1920 trials) over thirty consecutive sessions evenly spaced over the 6 weeks of training. The learning rate, quantifying how fast each participant converged to a relatively steady performance, varied significantly across participants (min = −0.96, max = −0.07; 25^th^ percentile = −0.36, median = −0.31, 75^th^ percentile = − 0.23), highlighting a substantial inter-individual difference in the sample tested (see Table S1 for the descriptive of the initial and final MT, and other parameters computed from the fitting procedure). Results and analyses of this section are further detailed in the supplementary material section 1.

### 2.2 Global and regional flexibility variation with ongoing practice

Based on the detected community assignments obtained for each participant and EEG session, we computed the flexibility of each of the 68 brain regions defined by the Desikan-Killiany anatomical atlas for each training intensity (EXT, MOD and MIN) and frequency band of interest. Flexibility was used to quantify the frequency at which brain regions change their community affiliation with time (i.e., across layers), normalized by the total possible changes. An assessment of this quantity at different levels of practice offers the advantage of quantifying the functional reorganization of specific brain regions throughout learning.

#### Quantification of global flexibility

We first sought to determine how brain network dynamics were configured as practice progresses, at a global level. To this end, changes in global flexibility (i.e., average of flexibility across brain regions) over the six weeks of training were evaluated with repeated measure ANOVAs (with training intensity -EXT, MOD and MIN- and laboratory session - 1 to 4 - as within-subject factors) for each frequency band (Figures 4a, 4c, 4e, and 4g). In all frequency bands, flexibility increased across sessions (*p* < .001) and significantly differed between EXT, MOD and MIN trials (*p* < .001), with highest values observed during the execution of the EXT sequences. Global flexibility increased more rapidly across scan sessions for sequences that were extensively practiced (interaction effect, *p* < .001). When data are sorted according to the number of trials performed (Figures 4b, 4d, 4f, and 4h; refer to Table S2 to see how cumulative practice trials were computed), they qualitatively replicate those shown in Bassett et al. (Bassett et al., 2013b) (see Figure 2c in their paper) where flexibility estimates were obtained from fMRI data recorded with the same experimental protocol used in this study. Consistently, we also observed an increasing number of communities with learning in all frequency bands (Figures S2e-h), although the quality of partitions in functional communities with increasing practice seems to increase in the theta and alpha bands and decrease in the beta and gamma bands (Figures S2a-d). Results are detailed in the supplementary material section 2.

**Figure 4:**
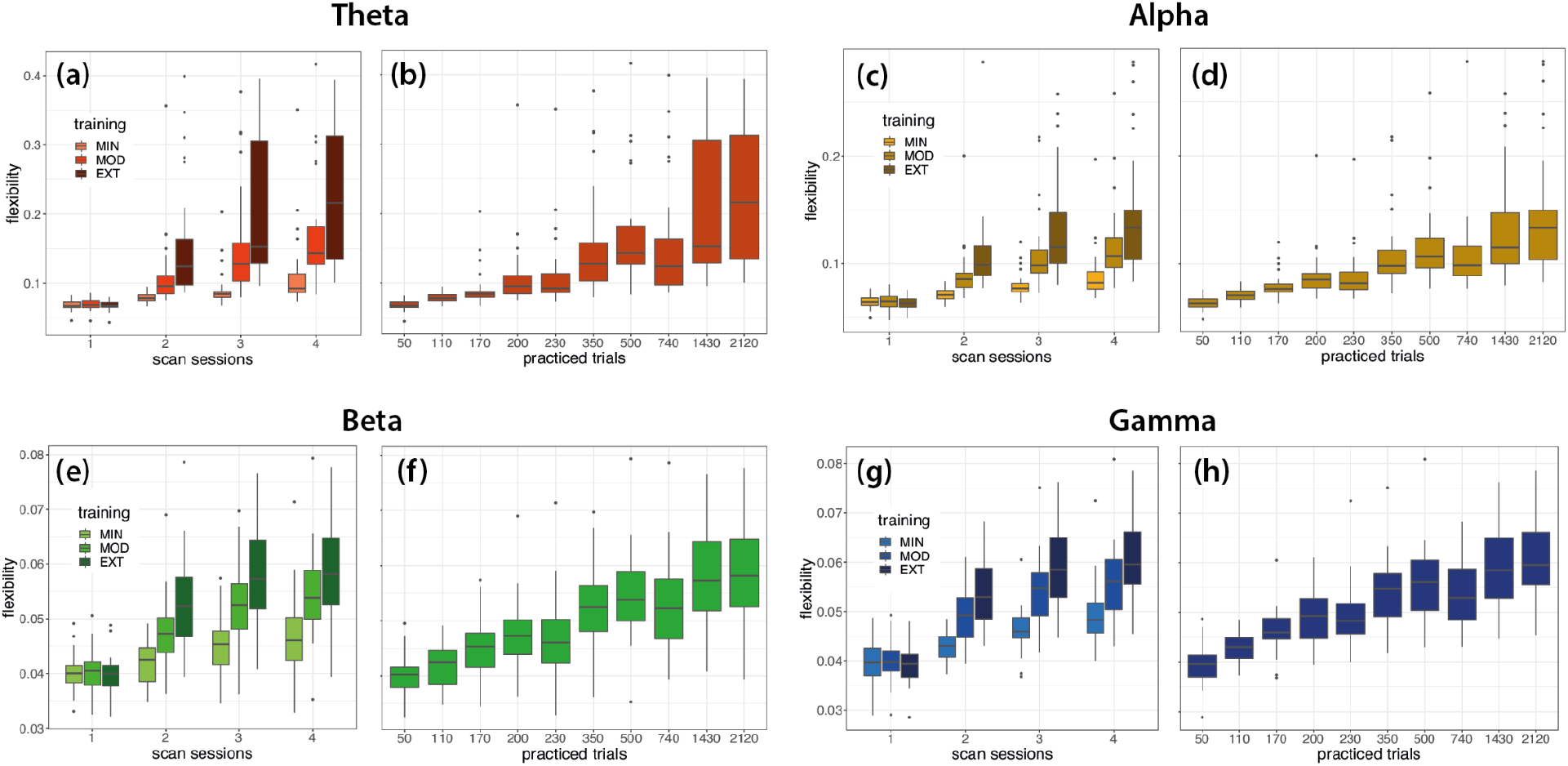
Global flexibility changes with learning. (a, c, e, g) Global flexibility, grouped by training intensity (EXT, MOD, and MIN), computed in the theta, alpha, beta and gamma bands from EEG data recorded at consecutive EEG sessions. (b, d, f, h) Global flexibility computed as a function of the number of trials completed after a given EEG session in the theta, alpha, beta and gamma bands. Lower and upper box boundaries represent the 25th and 75th percentiles (Q1 and Q3); the difference between Q1 and Q3 is the interquartile range (IQR); the horizontal line a inside the box represents the median of the distribution; the lower and upper extreme lines show Q1-1.5xIQR and Q3+1.5xIQR, respectively; finally, the filled gray circles are data points falling outside the lower and upper extreme lines.

#### Quantification of regional flexibility

To investigate the contribution of each brain region in increasing flexibility with practice, we contrasted flexibility values obtained from successive laboratory sessions for each frequency band. Significant differences were quantified with a one-tailed Wilcoxon test, testing the assumption that flexibility measured at a given session is bigger than flexibility at previous sessions. We included Bonferroni correction for multiple comparisons across brain regions. In Figure 5 we display regions with the most significant increase in flexibility between session 1 and 4.

**Figure 5:**
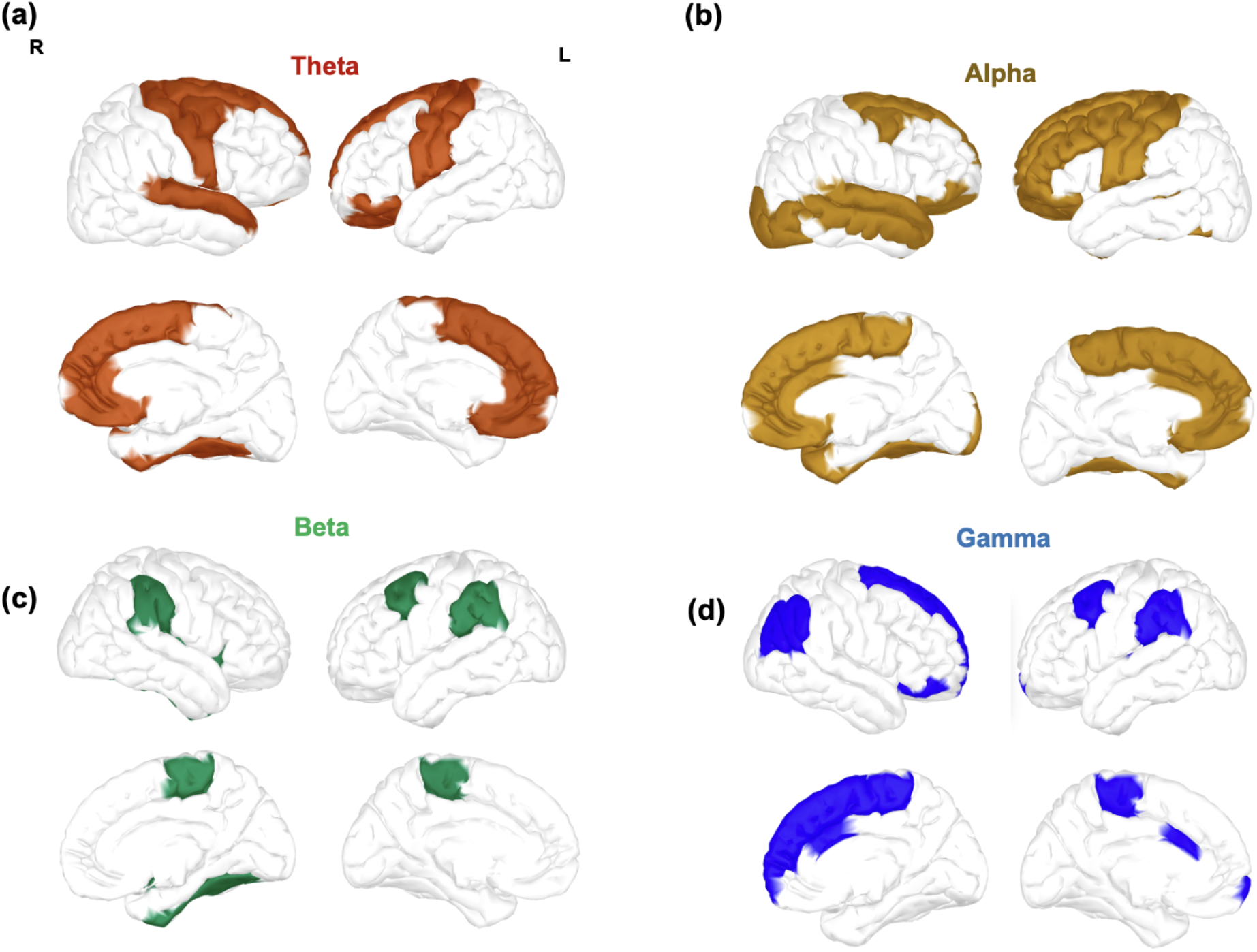
Brain regions displaying the most significant increase in flexibility between EEG recordings in session 1 and 4 for the (a) theta, (b) alpha, (c) beta, and (d) gamma frequency bands. Contrasts were assessed using a Wilcoxon test with Bonferroni correction, statistical results were divided in quintiles, and only statistically significant brain regions in the uppermost 2 quintiles are shown.

The theta and alpha bands (Figure 5a and 5b, respectively) were characterized by a similar increase of flexibility in prefrontal and central regions within default mode network (DMN) areas (rostral anterior cingulate, superior frontal, caudal middle frontal and superior temporal), in central regions within somatomotor network areas (precentral, postcentral and superior temporal), in prefrontal regions within limbic network areas (medial-orbitofrontal and rostral anterior cingulate), and in frontal regions over dorsal attention (superior frontal) and frontoparietal (caudal middle frontal) network areas. Specific to the alpha band (Figure 5b), we observed an increase in prefrontal regions within frontoparietal and ventral attention network areas (caudal anterior cingulate and left rostral middle frontal), over few more frontal regions within the DMN (caudal anterior cingulate and right middle temporal) and over posterior regions in visual network areas (right lateral occipital). Unlike for the theta and alpha bands, we did not observe a diffused significant increase in flexibility across brain regions in the beta frequency band (Figure 5c). Instead, flexibility specifically increased in the caudal middle frontal (covering areas within default mode and frontoparietal networks), in the parietal areas of the supra-marginal (within areas specific to dorsal, somatomotor and ventral networks), and in the paracentral within the somatomotor network. Finally, the gamma band (Figure 5d) was characterized by an increase in prefrontal (right superior frontal, caudal middle frontal and anterior cingulate, and pars orbitalis) and parietal (right inferior parietal) regions within the DMN areas, and in regions within areas of the frontoparietal (caudal middle frontal, caudal anterior cingulate and right inferior parietal), ventral (caudal anterior cingulate and left supra-marginal) and dorsal attention (right superior frontal, left supra-marginal and right inferior parietal), and somatomotor (left supra-marginal and paracentral) networks.

### 2.3 Functional correlates of performance

Next, we asked whether the flexibility of brain regions prior to extensive practice could be indicative of the individual learning profile observed during the following six weeks of training. To this end, we focused exclusively on the EEG signal recorded during the first EEG session. As described above, during this session participants practiced the six motor sequences for the very first time and the flexibility measures extracted during this recording could be indicative of the pre-learning functional state of each individual. To have consistent measures of flexibility across subjects, we thresholded the task trials length of each subject according to the length of the fastest recorded trial across subjects during the first session. As for the previous analysis, we built and analyzed the multilayer temporal networks through multilayer modularity optimization, and we used the computed flexibility as network diagnostics. We then correlated, for each region, the flexibility scores in each frequency band with the learning rate *K* obtained from practice during the following 6 weeks. Significant relationships - evaluated with Spearman’s rank correlations - were found in the beta frequency bands (Figure 6) in the left orbitofrontal (*ρ* = −0.50; *p*-value = 0.005) and rostral middle frontal (*ρ* = −0.50; *p*-value = 0.005) regions, and in the right parsorbitalis (*ρ* = −0.37; *p*-value = 0.044) and rostral anterior cingulate (*ρ* = −0.44; *p*-value = 0.014) regions. All the relationships remained significant after controlling for the initial and final MT, except for the right parsorbitalis region (Table S3).

**Figure 6:**
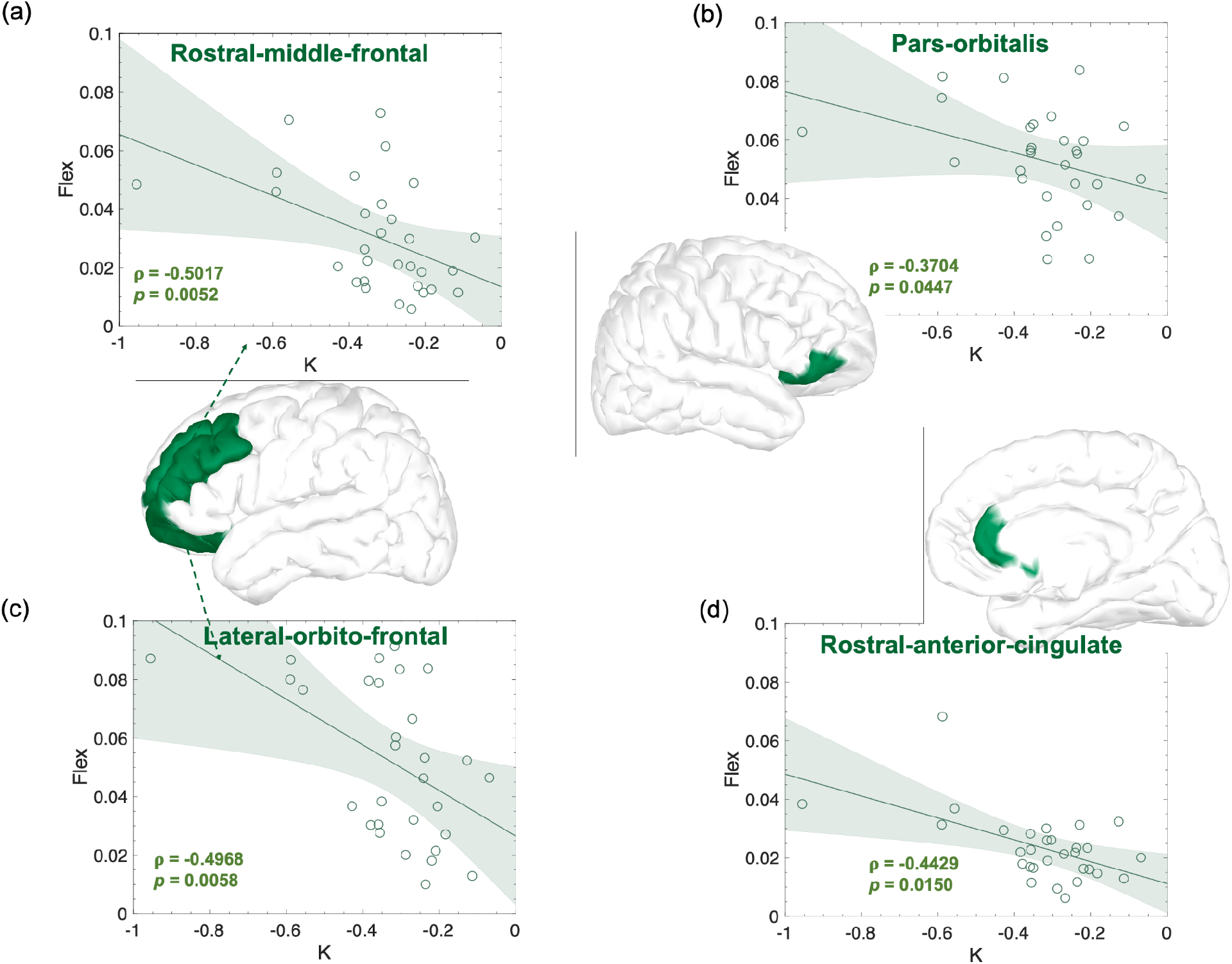
Significant correlations between regional flexibility in the beta frequency band computed from the EEG recorded in the first laboratory session and the learning rate K obtained from performance of the EXT sequences during home training sessions. Effect sizes (r and p) are also shown on the figure as well as the confidence intervals.

## 3. DISCUSSION

In this study, we aimed to assess whether extended practice of a new motor skill from visual cues was accompanied by frequency specific changes of brain network dynamics. We predicted a (i) global increase of cortical flexibility with learning. Specifically, we expected (ii) increased flexibility over prefrontal regions in the theta, alpha and beta frequency bands, and (iii) over sensorimotor and occipital regions in the alpha and beta bands. Lastly, (iv) specific to the theta, alpha and beta bands, we expected a positive relationship between participants’ prefrontal flexibility recorded before extended sequence practice and their learning rate from 6 weeks of training. Analyses in the gamma band were included to provide exploratory evidence of the dynamics of brain networks in these frequency ranges.

Thanks to the superior (sub-second) temporal accuracy of EEG compared to fMRI and the use of a longitudinal experimental setup (Figure 1 and 2), we were able to explore frequency-dependent aspects of learning by characterizing brain networks in specific EEG rhythms. We found that all participants improved performance of all sequences, with a strong inter-individual variability characterized by variable rates of improvement (Figure 3 & S1). We showed that the advancement of motor-skill learning through repeated and sustained sequence practice is accompanied by a robust reorganization of functional brain networks as shown by global flexibility increase in all studied frequency bands from theta to gamma ranges (Figure 4). As learning progresses, a considerable number of brain regions change more frequently their modular affiliation (Figure 5). Most of the increase was detected in the theta, alpha and gamma bands and, to a lesser extent, in the beta band. In particular, increases in numerous prefrontal and centro-parietal areas within DMN, frontoparietal and dorsal attention networks were observed in the theta, alpha and gamma bands, and over occipital regions within the visual network in the alpha band. Increases in prefrontal areas of the limbic network were specific to the theta and alpha bands. A consistent modulation of regions within the somatomotor network was instead revealed in all frequency bands. Finally, few regions within the ventral attention network increased their flexibility, mostly over frontal (alpha and gamma) and parietal (beta and gamma) areas. Finally, we have shown that individuals with highly flexible prefrontal regions in the beta spectrum at the onset of learning are those who learn more rapidly in the following 6 weeks of training (Figure 6).

The global increase of flexibility with practice is in line with previous fMRI studies (Bassett et al., 2013b, 2011; Reddy et al., 2018). Bassett and colleagues (Bassett et al., 2011) monitored performance on a simple motor skill task during three consecutive fMRI sessions and observed an increase in global flexibility during the early stages of learning followed by a relative decrease during the third recording. The lack of consistent flexibility increases as observed in our results could be explained by the three fMRI scans occurring at initial stages of learning. This did not allow an assessment of learning and flexibility over a longer period, as was instead possible in two subsequent fMRI studies (Bassett et al., 2013b; Reddy et al., 2018) assessing motor skill learning using the same longitudinal protocol used in our work. Consistent with our results (Figure 4b, d, f, and h), Bassett and colleagues (Bassett et al., 2013b) showed an increase in global flexibility (see Figure 2c in their work) with the number of trials practiced, which is consistent with an increased specificity of functional connectivity patterns with extended learning. On the same line, Reddy and colleagues. (Reddy et al., 2018) recently identified two “canonical” brain states made by sensorimotor and fronto-temporal subcortical regions during practice and observed that the rate of switching between these states constantly increased during the six weeks of training. Overall, the global flexibility increase observed in our study is coherent with a transition from greater behavioral adaptability towards more automatic performance with greater freedom of cognitive resources for other processes (Bassett et al., 2015; Shamloo and Helie, 2016).

As expected, we observed increased flexibility over prefrontal and limbic areas in the theta and alpha band, but we didn’t find a significant increase over these regions in the beta band. In terms of the brain areas involved and their dynamics with learning, our results are consistent with one of the core findings of Bassett and colleagues (Bassett et al., 2015) who highlighted a diffused decrease in the recruitment of prefrontal regions particularly relevant to the fronto-cingulate network. In line with their work, we observed variations of flexibility over hub regions (rostral anterior cingulate, caudal middle frontal, medial orbito-frontal and superior frontal) of cognitive control systems like the frontoparietal and cingulo-opercular networks (Elton and Gao, 2014). It was speculated (Bassett et al., 2015) that with learning these hubs become disengaged with the rest of the network as cognitive control is particularly critical during early skill acquisition (Hikosaka et al., 2002; Petersen et al., 1998). Our observed increase in flexibility supports this possibility. The fact that the increased flexibility with learning in these regions is specific to the theta and alpha rhythms is coherent with several lines of research. Our results are consistent with previous work highlighting a reduced coherence between prefrontal and central areas in both bands during movement planning and execution (Gentili et al., 2015) at later stages of learning. Similarly, our findings agree with those of Miraglia and colleagues (Miraglia et al., 2018) showing a global increase of modularity and transitivity measures after motor adaptation training in the theta and alpha band, and with those of Vecchio and colleagues (Vecchio et al., 2018) relating increased network segregation and integration with better learning in the alpha band. Importantly, Rilk et al. (Rilk et al., 2011) showed in a visuomotor tracking task that high tracking error was associated with enhanced frontocentral coupling in the alpha band, suggesting additional activation of a frontoparietal control network. Taken together, these are all evidence supporting the possibility that the release of specific hubs in the frontoparietal and cingulo-opercular networks (Bassett et al., 2015) occur within the theta and alpha frequency ranges. Finally, we observed unpredicted increases in few prefrontal regions of the right hemisphere in the gamma band. Although there is very little evidence for this frequency band, our findings seem coherent with a recently observed positive correlation between visuomotor tracking tasks difficulty and fronto-central connectivity (Aliakbaryhosseinabadi et al., 2021).

Coherent with our hypothesis, we found increased flexibility in the alpha band over somatomotor (pre and postcentral gyrus) and visual areas (lateral occipital regions). In the beta band, flexibility also increased over somatomotor areas but was only specific to paracentral and supra-marginal regions. Past research on how somatomotor areas change their activity during motor skill learning is divided into studies monitoring task-evoked BOLD activity (Dayan and Cohen, 2011) and connectivity (Bassett et al., 2015) during different phases of learning. Several studies showed a reduction in BOLD activity during the early stages of learning (Floyer-Lea and Matthews, 2005; Sakai et al., 1999) followed by an increase in activity during the late stages (Floyer-Lea and Matthews, 2005; Grafton et al., 2008; Honda et al., 1998). However, real insight into the dynamics of the undergoing process has been provided by connectivity studies and particularly by the work of Bassett and colleagues (Bassett et al., 2015). Consistent with the functional requirements of skill acquisition, they suggested that with extended practice motor and visual systems transition from being tightly integrated to operating as independent units. Our results of increased flexibility over somatomotor and occipital areas are coherent with these findings and go further suggesting that this late learning independence could be expressed by the increased frequency with which these regions change allegiance to functional communities. Only in the alpha band the modulations of flexibility covered areas related to a visuomotor network, whereas this was less evident in the beta band. Studies have shown that the coherence between motor and visual areas over centro-parietal and occipital regions in the alpha and beta band increases during the execution of visuomotor tasks (Classen et al., 1998; Erla et al., 2012), suggesting a functional link between these areas subserving sensorimotor integration. In addition, a higher coherence between these regions was associated with better performance during visuomotor tracking tasks (Rilk et al., 2011). However, these works say nothing about how connectivity varies with long-term learning but only provide information about the frequency ranges in which these modulations are supposed to occur. In contrast to these studies, we uncovered network adaptations across a continuum of learning and showed how the transition from more constrained toward more flexible dynamics happens in the alpha and, although less consistently, beta ranges. Finally, we observed unpredicted increases over somatomotor and centro-parietal areas in the theta and gamma bands. There is no specific evidence for the implication of somatomotor regions in the theta band, but our results are coherent with previous findings pointing to a reduced connectivity between frontal and central areas with ongoing visuomotor learning (Gentili et al., 2015). As for the gamma band, there is no concrete evidence highlighting a change in the dynamics of centro-parietal areas with visuomotor learning and our findings are not consistent with a recent study showing a gamma-related decrease in modularity following visuomotor adaptation training (Miraglia et al., 2018).

An important result of our work shows that some features of brain network dynamics measured in the very early stages of practice correlated very well with performance measured in independent sessions recorded over six weeks of practice. The most consistent results were found over prefrontal areas in the beta frequency band, including orbitofrontal, parsorbitalis and rostral middle frontal and anterior cingulate. Higher flexibility values were strongly correlated with faster reduction of movement time and thus, as suggested in previous work (Bassett et al., 2015; Wymbs and Grafton, 2015), faster learning. Previous fMRI studies have consistently shown that brain network flexibility correlates well with cognitive flexibility and learning (Bassett et al., 2013b, 2011; Braun et al., 2015; Reddy et al., 2018). In the context of simple motor skill learning, Bassett and colleagues (Bassett et al., 2011) showed that the amount of flexibility measured in one specific session correlated well with the amount of learning in the following practice session. Similarly, Bassett et al. (Bassett et al., 2013b) showed that the core-periphery geometry derived from each individual flexibility distribution during early learning correlated well with learning in the following ten days of training. Later, not directly addressing brain network flexibility, Bassett and colleagues (Bassett et al., 2015) showed that a consistent and gradual disengagement of prefrontal regions correlated with better learning. Our results are in line with these findings and go further by suggesting that early characteristics of prefrontal dynamics may be related to learning quality. We have noticed an absence of long-term longitudinal studies addressing (visuo)motor skill learning with EEG and lack of flexibility analysis that could be directly related to our work. However, our findings seem consistent with several lines of evidence from studies of visuomotor learning during periods of resting state. Consistent with our results and specific to the beta frequency range, Wu and colleagues (Wu et al., 2014) have shown that lower resting state connectivity between motor and left prefrontal areas was related to greater skill acquisition during subsequent training on a pursuit motor task. Although difficult to confirm without direct analysis, we speculate that lower connectivity between regions in these areas could translate into a more flexible modular composition over time and thus higher flexibility. In a similar study investigating arm reaching in a force mediated field, Faiman and colleagues. (Faiman et al., 2018) evidenced a positive relationship between resting state motor-prefrontal connectivity in the alpha band and performance. Their findings seem inconsistent with our results and with previous work (Wu et al., 2014). However, a point that is important to emphasize is the fact that the outcomes of Faiman and colleagues were obtained from analyses at scalp level, a factor that could explain their inconsistency. The implication of orbitofrontal and middle frontal areas in the beta frequency range is supported by evidence from other studies assessing brain dynamics during performance. A reduction in medial frontal beta-band activity (e.g., higher activation of the underlying area) during visual rotation was interpreted as the inhibition of automatic motor responses in favor of cognitively controlled movements (Jahani et al., 2020). We can speculate that a bias towards more cognitively controlled action may increase activity and connectivity between the areas involved, reducing temporal variability and thus flexibility. Finally, other studies have also shown relationships between brain dynamics at rest and visuomotor performance in other frequency bands (e.g., alpha (Manuel et al., 2018) and theta (Miraglia et al., 2018) bands), although these were not revealed by our study.

In conclusion, this work is the first attempt to characterize brain network dynamics in the context of motor skill learning by leveraging on a longitudinal design made up of several EEG recordings interspersed with long training periods. We confirmed previous evidence suggesting that with ongoing learning, cortical regions’ activity shifts from a constrained towards a less constrained, more flexible dynamics. In addition, we complemented traditional fMRI studies by providing access to the frequencies at which these changes occurred. Specifically, we have shown that the gradual disengagement of prefrontal regions highlighted in previous work is specific to cross-region interactions within frequencies from theta to alpha ranges, and that previous evidence on learning-related reduction of integration between sensorimotor and visual areas finds consistent support from the dynamics of these regions within the alpha frequency band. Importantly, we highlighted a relationship between the early dynamics of some prefrontal regions in the beta band and motor skill learning. This suggests some basic organizational principles and constraints of the brain already visible at the onset of learning.

### 3.1 Methodological considerations and limitations

Several important methodological and conceptual considerations are pertinent to this work. First, some details need to be discussed regarding the choice of experimental protocol used in this study, and the possibility that the observed behavioral and physiological outcomes are (at least in part) not directly linked to the experimental manipulation (in our case, practice of specific sequences and their learning). The experimental protocol used in this work is the same as that used by Bassett and colleagues (2013) and by other authors following their first publication (Bassett et al., 2015; Mattar et al., 2018; Reddy et al., 2018; Wymbs and Grafton, 2015). This choice was based on the compelling need to replicate recording procedures and analyses as consistently as possible with previous studies using the same experimental manipulation. Although the experimental protocol was not designed to include active control group or an additional experimental control condition, its design includes the presence of a within-participant factor through which the learning intensity is manipulated (on three levels: MIN, MOD and EXT) in order to control for non-specific familiarity effects due to the amount of time spent performing the experiment. For this reason, 2 sequences were practiced extensively, 2 occasionally, and 2 rarely throughout the training regime. This choice is justified by the evidence that the total amount of prior practice, rather than chronologic time, is the primary determinant of the magnitude and location of sequence-specific representations (Wymbs and Grafton, 2015). For this reason, there are valid reasons to assume that dynamics of changes in flexibility was largely related to learning and not necessarily due to the participant or the experimenter merely becoming accustomed to the experimental setting, or due to other changes relating to the passage of time or to general exposure to task practice. Indeed, the interaction effect between EEG session and training intensity level (MIN, MOD, and EXT) revealed a higher increase in flexibility with EEG sessions for the extensively practiced sequences than for the less practiced ones. During the last EEG scan, higher flexibility was observed during the practice of the EXT sequences compared to the MIN sequences, although the participants in the preceding weeks had received the same exposure to the task and experimental settings. However, the absence of an active group/control condition (designed in such a way as not to require learning - but with similar difficulties and manipulations of the training regime as in the experimental condition), poses the need to maintain relative caution regarding the effects of the experimental intervention presented in this and previous studies. To definitively exclude the possibility that the observed effects are due to factors external to learning, future studies should consider the inclusion of an additional control condition/group (e.g., via the deployment of simple motor tasks). Nevertheless, we can already anticipate the difficulty in reaching such a compromise given that the additional control group / condition should be designed to solicit cognitive-motor processes comparably to the experimental condition, as well as match in difficulty and motivation for the participants, all of which are extremely difficult factors to control.

Second, a further point of discussion relates to the use of flexibility as a quantitative measure to describe functional brain dynamics and its link with learning. The use of this parameter allowed us to compare the results obtained in this study with those obtained in previous fMRI studies with similar protocols. However, an inherent limitation of this measure is that it does not provide the level of detail required to infer the functional mechanisms that generate the observed variations in flexibility. For instance, the same flexibility value can be generated by a region changing, with a certain frequency, affiliation to only two or more communities. This is complicated by the fact that the size and spatial location of each community obtained through modularity maximization is dynamic and therefore varies over time. Future studies could focus effort on research and implementation of quantitative measures possibly complementary to the approach followed in this work such as metrics related to network segregation (such as clustering coefficient and recruitment), integration (such as participation coefficient) and hubness (such as betweenness centrality). Morever, measures such as global and nodal efficiency could also be implemented to test a plausible hypothesis that learning favors faster communication and less effort in transferring information. A final consideration must be made with respect to the scale resolution of analysis presented in this work, as this study focused on dynamic variations occurring at a larger time scale corresponding to the time period of task execution. However, the available dataset would in principle allow adapting existing analysis procedures to enable a finer scale analysis of the dynamics occurring at the scale of the individual trial. This possibility might uncover additional features that would enhance understanding of functional network-based predictors of learning phenomena.

Third, with respect to the computation of functional connectivity to build multilayer modularity tensors, it is important to note that the dynamic connectivity matrices computed in each trial (windowed PLVs) were not used in our work other than to calculate the values of the static PLVs in each trial. It is in fact the latter that were used as inputs for the construction of the multilayer network tensors for community detection. An equivalent and computationally more advantageous alternative would have been to calculate the static PLVs directly using the entire signal within each trial, thus avoiding the intermediate step involving the calculation of the wind owed PLVs. Nevertheless, our choice to go through the additional step of calculating the windowed PLVs has the advantage that the database that we created could be used as a basis for more detailed analyses (e.g., for questions specific to temporal dynamics at the level of the individual trial) in future studies, with the possibility of comparing in a more consistent manner future findings with the results obtained in this work.

Fourth, with respect to the algorithm used to calculate global and regional flexibility, we replicated in this work the methodological steps described in previous works that used the same or similar protocols to the one used in (Bassett et al., 2013b, 2011; Betzel et al., 2017). Specifically, as described in section 4.5, the flexibility at the subject level was computed by averaging the flexibility values obtained from different partitions obtained by multiple maximizations of the multilayer modularity function (this was done for each session, training intensity level, and frequency band). This approach made it possible to limit near-degeneracies in the modularity landscape and to obtain flexibility estimates that were consistent and comparable with those obtained in the previous studies cited above. However, several valid alternatives may exist to the procedure implemented here. These include the possibility of using a consensus partition from the realizations obtained from the maximization process. In this sense, a recent study has proposed the use of a generative modeling approach called weighted stochastic clock models (WSBM) as an effective tool to describe different types of community structure in brain networks that goes beyond simple modularity (Faskowitz et al., 2018). Consensus partitions and flexibility measures can be estimated using this method. This method was successfully applied to study changes in the human connectome that occur across the lifespan, and has indeed great potential to be implemented on longitudinal datasets like ours to identify how community structure regimes relate to aspects of behavior and cognition, such as learning.

Fifth, our sample of young adults has, out of 30 participants, 25 female volunteers. Testing mainly females makes the generalization of our results slightly less robust, although there are premises in the literature suggesting no advantage for either female or male adults. First, previous studies using the same task as ours reported no gender differences in the variables studied (Bassett et al., 2015, 2013b, 2011; Mattar et al., 2018; Reddy et al., 2018; Wymbs and Grafton, 2015). Second, and with respect to various contexts of motor performance, evidence in the literature has reported an advantage for women in fine-motor tasks (e.g., pegboard tasks, handwriting: Berninger et al., 2008; Bornstein, 1986), an advantage for men in speed execution but not in accuracy of purely motor tasks (e.g., finger tapping: Gur et al., 2010; Hausmann et al., 2004; Nicholson and Kimura, 1996; Ruff and Parker, 1993), but no clear advantage for either male or female adults in performance of sensorimotor tasks (e.g., Gur et al., 2012, 2010). In addition, evidence on performance of professional musicians during sequential motor skill tasks like our DSP task (e.g., Kincaid, Anthony E. et al., 2002) did not reveal any gender differences in either professional musicians or controls. Finally, control on gender by introducing it as a between-subject factor in our analyses of variance did not reveal any significant difference in movement time and learning parameter estimates. However, these analyses should be interpreted considering the small sample and the unequal number of subjects in each factor’s level.

Finally, we explored the possible confounders related to two additional factors - EEG signal strength and task design - and were able to put some elements into perspective (see supplementary material, section 4). First, we have shown that the flexibility results obtained through phase-related analyses of the EEG signal are independent of the amplitude content of the signal itself (section 4.1). Second, we have highlighted that flexibility was correlated with variables related to the implicit structure of the task (e.g., trial length). We have suggested that properties of multilayer networks and flexibility may have a common origin due to reduced signal length with learning, although it was impossible to infer a causal role between those variables. In this regard, we have provided additional analysis (sections 4.2.1 & 4.2.2) to take these variables into account but are aware of the multiple limitations of this approach. Amongst these, the most critical is the loss of a significant amount of EEG signal due to the proposed thresholding procedure to resize the trials to the same length. Nevertheless, despite these limitations, the results obtained are in partial overlap with the main findings, which is reassuring considering previous work on this topic (Bassett et al., 2015, 2013b; Reddy et al., 2018). However, we argue that the elements raised here must be seriously taken into consideration as they represent an interesting opening for future work: because of the difficulty to disentangle mechanisms that compete with the type of tasks typically developed and used in the learning paradigms, we flag as a priority the development and use of new experimental protocols that are tailored to the methodology used.

## 4 MATERIALS AND METHODS

### 4.1 Participants

Thirty healthy volunteers (25 females; mean age = 22.2 years, SD = 6.2 years, range 18 to 50 years) gave written informed consent to participate in the experiment and were compensated with credits for their participation. All participants reported normal or corrected-to-normal vision acuity and a right-hand preference (84 ± 22) (Oldfield, 1971). Research was approved by the Cantonal Ethics Committee for Human Research (Vaud, Switzerland; protocol N°2019/02345) and was in accordance with the code of ethics of the World Medical Association (Declaration of Helsinki) for experiments involving human subjects in research.

### 4.2 Experimental setup and procedure

We used the same experimental protocol developed by Bassett et al. (2013b) and previously employed in a large variety of studies (Bassett et al., 2015; Mattar et al., 2018; Wymbs and Grafton, 2015). The protocol (Figure 1a) is a longitudinal study consisting of home and laboratory sessions. Participants practiced 30 home training sessions of a discrete sequence-production task (DSP) over 42 days at a rate of 10 sessions every 14 days. In addition, they were tested on the same DSP task during 4 laboratory sessions where EEG was recorded. The first laboratory session was planned at the beginning of the study, and the remaining three every 14 days after 10 consecutive home sessions.

During home sessions (Figure 1b) participants practiced a total of 150 DSP task trials, consisting of two sequences (EXT1 & EXT2) each presented for 64 trials (extensively trained sequences; EXT), two sequences (MOD1 & MOD2) for 10 trials (moderately trained sequences; MOD), and two additional sequences (MIN1 & MIN2) for 1 trial (minimally trained sequences; MIN). Each sequence was made of 10 sequential visual displays. Each visual display contained a horizontal array of five square stimuli, each corresponding to one finger (from left to right, the thumb and the pinky were placed on the keyboard’s “spacebar” and on the letter “L” and corresponded to the leftmost and rightmost square, respectively; the index, middle and ring fingers on the letters “H”, “J” and “K”). In each display, one of the five squares had a red outline, indicating which key to press. Immediately after pressing the correct key, the display changed, and another square switched its outline to red. Participants could only advance the sequence by pressing the correct keypress and had unlimited time to complete it as accurately and quickly as possible. In case of error, an error message appeared on the screen for 100 ms and the sequence was paused at the error and restarted upon the appropriate key press. Each sequence was conceived such that each of the five square stimuli switched twice its outline to red, and to avoid immediate repetitions (e.g., “1-1”) and regularities (e.g., “4-5-4” or “4-3-2”) across consecutive visual displays. Each trial started with the presentation of a cue at the center of the screen for 500 ms (red and blue circle for the EXT sequences, green and yellow triangles for the MOD sequences, white and black stars for the MIN sequences), followed by the first of the ten visual displays of the sequence. The end of the sequence was followed by a 200 ms screen-centered fixation cross, and an inter-trial blank screen varying in duration between 500 and 1500 ms. For every block of 10 trials, feedback about the number of error-free sequences and mean time to complete the error-free sequences was presented for 3000 ms right after the fixation cross. In these trials, the feedback was followed by a random inter-trial interval lasting between 0 and 1000 ms. EXT, MOD and MIN trials were randomly distributed throughout the task.

During EEG sessions (Figure 1b) participants practiced EXT, MOD and MIN sequences for 50 trials each, for a total of 300 trials. The trial design was the same as in the home sessions. However, unlike the home sessions, the trials were not presented randomly but evenly distributed over 5 epochs of 60 trials each (i.e., each epoch contained the same number of EXT, MOD and MIN trials). Each epoch contained 2 blocks of 10 EXT trials (5 trials for each of the 2 EXT sequences), 2 blocks of MOD trials (5 trials for each of the 2 MOD sequences), and 2 blocks of MIN trials (5 trials for each of the 2 MIN sequences). Trials were randomized within each block and blocks randomized within each epoch.

The DSP task practiced during home sessions was coded with PsychoPy (Peirce et al., 2019) in JavaScript and was made available in the form of a web-based task that participants could access from their personal computer. Participants were asked not to perform close sessions and to keep the same daily schedule and environment as much as possible for home practice. The DSP task practiced during laboratory sessions was coded with Psychopy and run on the laboratory computers. Stimuli were synchronized with the EEG hardware by means of TTL-triggers marking the EEG signal with task-related events (e.g., visual displays onsets). During these sessions, participants were seated in a quiet room sheltered from electromagnetic disturbances (Faraday cage). Stimulus size and participant’s distance from the screen were adjusted so that the 5 square stimuli of the DSP task covered a horizontal length of 7° visual angle. To limit EEG artifacts due to head movements, the head of participants was placed on a chin rest.

### 4.3 Behavioral estimates of learning

Previous works using the same experimental procedure (Bassett et al., 2015, 2013b; Mattar et al., 2018; Wymbs and Grafton, 2015), have quantified behavioral learning with movement time (MT), defined as the time that elapses between the first and last button press during the execution of a sequence. The decrease in MT with practice has often been used to quantify learning (Crossman, 1959; Stratton et al., 2007). Various functional forms have been used to model this decrease and the most robust choices have often been made in favor of (variant of) exponential functions (Heathcote et al., 2000). To quantify the decrease of MT with practice, we used in this work an approach similar to previous studies which modeled the MTs associated to EXT sequences using a double exponential function (Bassett et al., 2015, 2013b; Mattar et al., 2018; Wymbs and Grafton, 2015). This choice was motivated by the vastly superior number of practiced trials in EXT sequences. First, MTs were calculated for each sequence performed in the home sessions during the 42 days of practice. The MT was computed from the first to the last button press of the ten displays. In case of error, the time spent visualizing the error display was not included into the computation of MT. The MTs of the two EXT sequences were then averaged together for each of the 30 home-based sessions, resulting in 30 MT values for each participant. These data were then fitted with a double exponential function of the form (1):

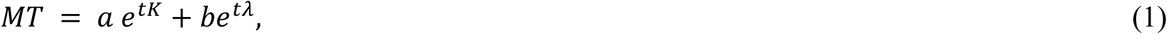

where *t* is time, *K* ∈ (-∞,0] is the exponential drop-off parameter describing the fast rate of improvement (typically occurring during the first days of practice), λ ∈ (-∞,0] is the exponential drop-off parameter describing the slow, sustained rate of improvement, and *a* and *b* ∈ [0,+∞) are constants. The parameter *K*, which is independent of the initial and final individual’s performance, was used as an indicator of the speed of learning as in previous works (Bassett et al., 2015; Dayan and Cohen, 2011): curves with high absolute values of *K* are characterized by a higher improvement of MT per unit of time (i.e, decrease more rapidly) than those described by smaller values. In addition to the learning rate *K* it was also computed, for each participant, the initial and final MT (average MT computed from EXT sequences in the first and last home training session).

### 4.4 EEG recording, preprocessing and functional connectivity estimation

Continuous EEG was obtained with a Biosemi ActiveTwo system using 128 active electrodes placed at the international radial ABC system location and two pairs of bipolar electrodes recorded EOG in both vertical and horizontal directions. Two additional electrodes (active CMS: common mode sense, and passive DRL: driven right leg) were used as reference and ground to compose a feedback loop for amplifier reference. Data sampling was set at 1024 Hz with 24-bit A/D conversion. EEG data were acquired at each EEG session during performance on the DSP task, and during two 5-min resting state periods occurring at the beginning and at the end of the experimental session (the reader should refer to Figure 2 for a visual representation of the methodological steps described in the following paragraphs).

We first band-pass filtered the EEG signals (1-45 Hz) and we extracted trial epochs from the onset of the first display to the offset of the last sequence display, with a 500ms pre-stimulus baseline. We removed eye blinks using the EOG regression method (i.e. use the EOG channels to clean the 128 EEG channels) (Parra et al., 2005). Channels with high standard deviation and flat waveform were identified as bad channels and interpolated using the spherical spline method as implemented in eeglab (Delorme and Makeig, 2004). Signals were then re-referenced to an average reference montage. We then computed several quality check parameters to rank the preprocessed epochs from high to lower quality, namely the Overall High Amplitude, the Time High Variance and the Channel High Variance. All these metrics are described in Automagic (Pedroni et al., 2019). This was used as an additional step to remove bad trials with very low-quality metrics. We finally performed an additional visual inspection of the trials based on topographies displayed at the max of the Global Field Power (GFP) within each trial.

After preprocessing the EEGs, the functional connectivity matrices were estimated in each trial using the EEG source-connectivity method (Hassan and Wendling, 2018; Schoffelen and Gross, 2009). First the dynamics of cortical brain sources were reconstructed by solving the inverse problem. To do so, EEG channel locations and MRI template (ICBM152) were co-registered, and using the OpenMEEG toolbox (Gramfort et al., 2010), a realistic head model based on the Boundary Element Method (BEM) with three layers (scalp, skull and brain) was built. The weighted minimum norm estimate (wMNE) was used to estimate the brain sources on a surface mesh of 15002 vertices. The regularization parameter is set to 0.3 in our study. The noise covariance matrix was computed from a 500 ms pre-stimulus baseline. In this work, we used the Matlab function implemented in Brainstorm toolbox (Tadel et al., 2019) to compute wMNE, with the signal to noise ratio set to 3 and depth weighting value to 0.5 (default values). The regional time series of the 68 cortical regions of interest (ROI) of the Desikan-Killiany atlas were computed by averaging the activity of sources included in each ROI (Figure 2a).

Afterwards, we filtered the regional time series in different EEG frequency bands: Theta: 4-8 Hz; Alpha: 8-12 Hz; Beta: 12-28 Hz; Gamma: 28-45 Hz. For each frequency band, we computed functional connectivity using the Phase Locking Value (PLV), a measure for assessing phase synchrony between two signals in a particular frequency band, formed from estimates of the instantaneous phase of the signal. More specifically, we inferred connectivity at each trial by applying a sliding window approach where connectivity was computed within each temporal window (dynamic or windowed PLVs) and then averaged across windows to obtain a single connectivity matrix (static PLV). This was done by taking into account the recommendations of Lachaux et al. (1999) concerning the choice of smallest window length to guarantee a sufficient number of cycles per window at the given frequency band. This minimum window length equals 6 divided by the central frequency of each frequency band, where the number 6 corresponds to the smallest number of cycles recommended by Lachaux et al. (1999). We therefore ended up, for each frequency band, with one static PLV for each trial (Figure 2b). These static PLVs from the task trials of each session were used in the following analyses as inputs for the construction of the multilayer network tensors for community detection (Figure 2c).

### 4.5 Construction of dynamic networks, community detection and flexibility estimates

We characterized brain networks in terms of global cohesive functional structures that could capture relevant dynamics underlying motor skill learning occurring at time scales of several weeks of practice. We first constructed multilayer network tensors **A** (Mucha et al., 2010), representing time-dependent networks (Figure 2c), by temporally combining the static connectivity matrices (static PLVs) from *L* consecutive trials of a given sequence type (EXT, MOD, and MIN). This was done, in each frequency band, for each participant and EEG session. No thresholding was applied to the static PLVs. Starting from these multilayer network tensors, we then identified functional communities (or modules) composed of brain regions exhibiting similar functional activity through time. To this end, we performed community detection (Fortunato, 2010; Porter et al., 2009) on each multilayer network tensor in order to identify groups of brain regions that have a strong connection to each other (and preserved across temporal layers) with respect to brain regions assigned to other groups (Figure 2d).

To obtain functional communities we opted for the method of modularity maximization (Mucha et al., 2010) through the use of the Louvain-like locally greedy algorithm (Blondel et al., 2008), recently adapted and used in the context of studies linking multilayer networks to motor skill learning (Bassett et al., 2015, 2013b, 2013a, 2011). Modularity maximization consists in the optimization of a multilayer modularity quality function *Q* (equation 2) defined as in Mucha et al. (Mucha et al., 2010):

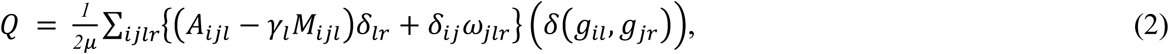

where *μ* is the sum of edge weights across nodes and layers, *δ*_ij_ equals 1 when *i* = *j* and equals 0 otherwise, and δ_lr_ equals 1 when *l* = *r* and equals 0 otherwise. A_ijl_ is an element of the multilayer network tensor **A**, representing the strength of the connectivity edge between node *i* and *j* in layer *l*, while M_ijl_ is the strength expected from a Newman-Girvan null model (Bassett et al., 2013a). γ_l_ is the structural resolution parameter of layer *l* (intralayer coupling parameter), the quantity *g*_*il*_ and *g*_*jr*_ give the community assignment of node *i* and *j* in layer *l* and *r*, respectively (δ(*g*_*il*_, *g*_*jr*_) equals 1 when *g*_*il*_ = *g*_*jr*_, 0 otherwise). ω_jlr_ is the interlayer coupling parameter (often referred as temporal resolution parameter) representing the connection strength from node *j* in layer *l* to node *j* in layer *r*.

We performed multilayer modularity maximization using the generalized Louvain package in MATLAB implemented with the Modularity Probability Method (MPM) algorithm (see (Bazzi et al., 2016; Yang et al., 2021) for details on how the MPM algorithm is implemented to detect and merge communities), with γ = ω = 1 (Bassett et al., 2015, 2013a, 2011). To overcome near-degeneracies in the modularity landscape which is expressed in a different community partition at each run of the algorithm optimizing *Q*, we avoid focusing on a single algorithmic run (Bassett et al., 2013a). Instead, following the approach proposed in previous works that have also used flexibility as a quantitative metric for their analyses (Bassett et al., 2013b; Betzel et al., 2017), we optimized each multilayer modularity quality function *Q* 100 times and computed regional and global network flexibility for each algorithmic run and subsequently averaged over the 100 runs. We computed regional network flexibility for each of the brain regions defined by the Desikan-Killiany template (Figure 2e). It is an indicator of the stability of functional communities, derived from the optimization of the quality function *Q*, across temporal layers. Specifically, it quantifies the fraction of times that a specific brain region changes assignment to a community in successive temporal layers, normalized on the total number of possible changes. A value close to 0 indicates great consistency in community assignment over time (i.e., over the course of a scan session for a specific sequence type - EXT, MOD or MIN), while a value close to 1 indicates great variability. Global network flexibility was computed by averaging regional flexibility across all brain regions. Regions in which significant changes in flexibility are observed are named using the Desikan-Killiany template. Throughout the text, an indication of the possible brain networks with which they are associated is also provided (Thomas Yeo et al., 2011).

### 4.6 Statistical modeling

Using a two-way repeated-measures analysis of variance (ANOVA), we tested the effect of laboratory sessions (session 1 to 4) and training intensity (MIN, MOD, EXT) on the MT (section 2.1 & supplementary section 1), the global flexibility estimates (section 2.2 & supplementary section 2 - quantification of global flexibility) and power estimates (supplementary section 4 - relationship between EEG signal power and flexibility estimates) in each frequency band. We applied a correction of the *F*-values in case of violation of sphericity assumption (Mauchly’s test) using a Greenhouse-Geisser (if Greenhouse-Geisser epsilon < 0.75) or a Huynh-Feldt (if Greenhouse-Geisser epsilon >= 0.75) correction. We interpreted significant interactions using pairwise comparisons and applied Bonferroni corrections to address the problem of multiple comparisons. To identify the cortical regions showing significant regional flexibility values, we used a one-tailed Wilcoxon test and ran Bonferroni and False Discovery Rate (FDR) corrections to address the problem of multiple comparisons. Finally, we used Spearman’s rank correlation as a measure of statistical relationship between flexibility and behavior (section 2.3), power estimates (supplementary section 4) and variables used to characterize task design (supplementary section 4). This nonparametric statistic measures the monotonic relationship between two variables without a requirement for linearity. ANOVAs were performed with the Jamovi project (2021)(“The jamovi project (2021). Jamovi (Version 1.0.7.0) [Computer Software]. Retrieved from https://www.jamovi.org.,” n.d.); all other analyses were performed in MathWorks MATLAB R2020b implementing custom code.

## DATA AND CODE

Data and code are made available on the open access repository Zenodo (Ruggeri, Paolo, 2022).

