## Supplementary File text for "Dynamic rewiring of electrophysiological brain networks during learning"

#### Supplementary text

Paolo Ruggeri[1], Jenifer Miehlbradt[1], Aya Kabbara[2,4] and Mahmoud Hassan[3,4]

[1] Brain Electrophysiology Attention Movement Laboratory, Institute of Psychology, University of Lausanne, Switzerland.

[2] Lebanese Association for Scientific Research, Tripoli, Lebanon

[3] School of engineering, University of Reykjavik, Reykjavik, Iceland.

[4] MINDig, F-35000 Rennes, France

##### Supplementary section 1: Behavioral performance

Repeated measure ANOVAs with training intensity (EXT, MOD and MIN) and EEG session (session 1, session 2, session 3 and session 4) as within-subject factors were applied to the MT recorded during EEG sessions (Figures 3a-b). We found a main session effect ($F(3, 87) = 465.9$, $p < .001$, $\eta^2_p = 0.941$), and bonferroni-corrected post-hoc comparisons revealed that MT significantly decreased across sessions ($p < .001$; S1: $M = 4.66$ s, $SE = 0.19$ s; S2: $M = 2.44$ s, $SE = 0.11$ s; S3: $M = 1.94$ s, $SE = 0.10$ s; S4: $M = 1.65$ s, $SE = 0.09$ s). A main effect of training intensity was also observed ($F(2, 58) = 359.6$, $p < .001$, $\eta^2_p = 0.925$). Bonferroni-corrected post-hoc comparisons revealed significant differences in the MT to execute EXT, MOD and MIN sequences, with lowest values to execute EXT sequences ($M = 2.22$ s, $SE = 0.11$ s), followed by MOD ($M = 2.54$ s, $SE = 0.13$ s) and MIN ($M = 3.26$ s, $SE = 0.12$ s) sequences. There was a significant interaction between training intensity and session (F(6, 174) = 95.8, p < .001, η2p= 0.768), and bonferroni post-hoc tests revealed that MT decreased more rapidly across scan sessions for sequences that were extensively practiced during home-based sessions as compared to less practiced ones ($\Delta(S2\text{-}S1)_{EXT} = -3.02$ s, SE = 0.12 s, t = 24.921, p < .001; $\Delta(S2\text{-}S1)_{MOD}$ = -

2.31 s, SE = 0.13 s, t = 17.588, $p < .001$; Δ(S2-S1)MIN = -1.30 s, SE = 0.11 s, t = 11.469, $p < .001$).

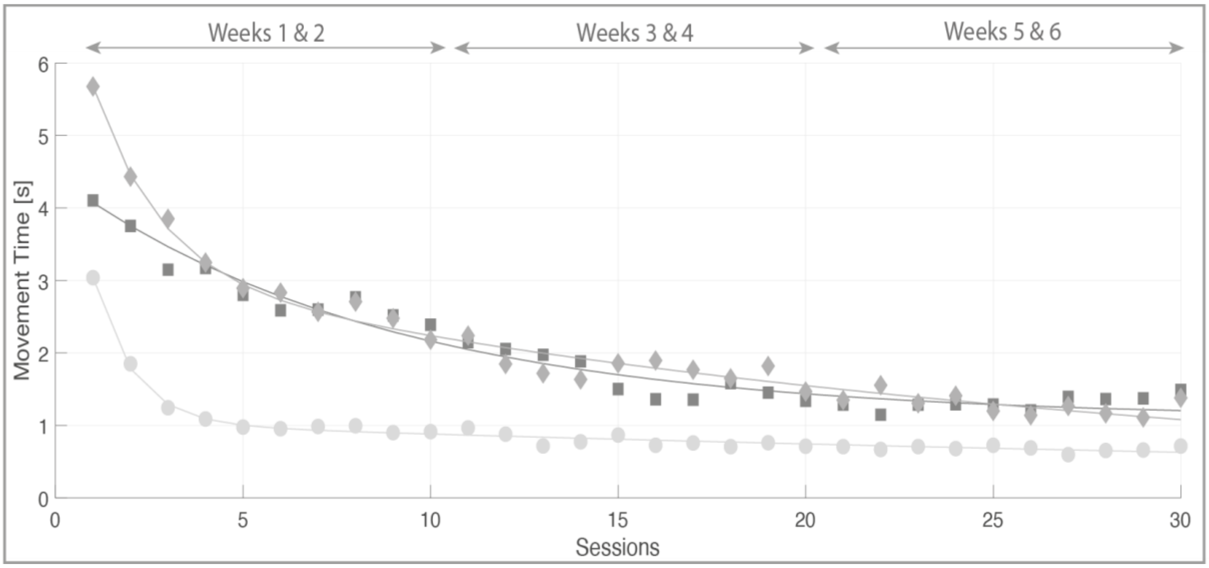


**Figure S1: MT for the EXT sequences recorded during home training sessions and the corresponding double-exponential fit. MT fits are displayed for three representative participants with different learning rate *K*: light gray curve marked circles (fast decrease of MT with practice; *K* = -0.96), medium gray curve marked diamonds (moderate decrease of MT with practice; *K* = -0.56) and dark gray curve marked squares (slow decrease of MT with practice; *K* = -0.11).**

In table S1 we report the descriptive of the parameter estimates obtained from the double exponential fitting procedure applied to the MT recorded during the home sessions. Relative to the MT, two additional quantities were also computed: the initial MT obtained from the average MT in the first home session, and the final MT from the average MT from the last two home sessions.

| Parameter | MIN | MAX | MEDIAN | 25th percentile | 75th percentile |
|---|---|---|---|---|---|
| **a [sec]** | 1.56 | 5.24 | 3.27 | 2.55 | 4.11 |
| **K** | -0.96 | -0.07 | -0.31 | -0.36 | -0.23 |
| **b [sec]** | 0.54 | 3.50 | 1.43 | 1.10 | 2.04 |
| **$\lambda$** | -0.036 | 0 | -0.009 | -0.014 | -0.001 |
| **MT init. [sec]** | 1.93 | 5.68 | 4.06 | 3.50 | 4.65 |
| **MT fin. [sec]** | 0.57 | 2.58 | 1.20 | 1.07 | 1.47 |

**Table S1: Minimum, maximum, and percentile values (25th, median, and 75th) of the double exponential fitting parameters (a, K, b, and $\lambda$) and of the initial and final MT distributions computed from home session recordings of 30 participants during practice of EXT sequences.**

Gender differences were checked in our data by introducing gender as a between-subject factor in the analyses of variance. The analyses did not reveal any significant gender difference in movement time and learning parameter estimates. These results should be understood considering the small sample and the unequal number of subjects in each factor's level.

#### Supplementary section 2: Global and regional flexibility variation with ongoing practice

##### Quantification of global flexibility

We evaluated changes in global flexibility (i.e., average of flexibility across brain regions) over the six weeks of motor skills practice, for each frequency band (Figure 4). Repeated measure ANOVAs with training intensity (EXT, MOD and MIN) and EEG session (session 1, session 2, session 3 and session 4) as within-subject factors were applied to the global flexibility estimates for each frequency band (Figures 4a, 3c, 3e, and 3g). A main effect of session was found in the theta ($F(1.58, 45.82) = 54.9$, $p < .001$, $\eta^2_p = 0.655$), alpha ($F(1.66, 48.27) = 59.2$, $p < .001$, $\eta^2_p = 0.671$), beta ($F(2.15, 62.24) = 89.6$, $p < .001$, $\eta^2_p = 0.756$), and gamma ($F(2, 58) = 147.9$, $p < .001$, $\eta^2_p = 0.836$) band. Bonferroni-corrected post-hoc comparisons revealed that global flexibility significantly increased across sessions in all frequency bands except for the increase between session 3 and 4 in the beta band ($p = .160$). A main effect of training intensity was observed in the theta ($F(1.34, 39) = 53.1$, $p < .001$, $\eta^2_p = 0.647$), alpha ($F(1.22, 35.28) = 49.7$, $p < .001$, $\eta^2_p = 0.632$), beta ($F(1.27, 36.94) = 146.4$, $p < .001$, $\eta^2_p = 0.835$) and gamma band ($F(1.40, 40.54) = 184.3$, $p < .001$, $\eta^2_p = 0.864$). Bonferroni-corrected post-hoc comparisons in all frequency bands revealed that global flexibility significantly differed between EXT, MOD and MIN trials, with highest values observed during the execution of the EXT sequences, followed by MOD sequences and smallest values for the MIN sequences. A significant interaction between training intensity and session was also observed for all frequency bands: theta ($F(3.46, 100.35) = 24.2$, $p < .001$, $\eta^2_p = 0.455$); alpha ($F(2.96, 85.88) = 26.1$, $p < .001$, $\eta^2_p = 0.474$); beta (F(3.87, 112.09) = 49.7, p < .001, η2p= 0.631); gamma (F(4.18, 121.15) = 56.6, p < .001, η2p= 0.661). Bonferroni post-hoc tests revealed that, in each frequency band, global flexibility increased more rapidly across scan sessions for sequences that were extensively practiced during home-based sessions as compared to less practiced ones (e.g., for the theta band: $\Delta(S2\text{-}S1)EXT = 0.089$, SE = 0.015, t = 6.079, $p < .001$; $\Delta(S2\text{-}S1)MOD = 0.042$, SE = 0.009, t = 4.688, $p = .004$; $\Delta(S2\text{-}S1)MIN = 0.010$, SE = 0.001, $t = 8.136$, $p < .001$). In Figures 4b-d-f-h the same results displayed in Figures 4a-c-e-g are shown sorted according to the number of trials performed (refer to Table S2 to see how cumulative practice trials were computed). These results qualitatively replicate those shown in Bassett et al. (2013) (see Figure 2c in their paper) where flexibility estimates were obtained from fMRI data recorded with the same experimental protocol used in this study. According to this pioneering study by Bassett et al. (2013), the systematic increase in global flexibility with practice observed in our study could be consistent with an increased specificity of functional connectivity patterns with extended learning.

In Figure S2 we show the variation of the multilayer modularity quality function *Q* (i.e., its maximum value resulting from its optimization as provided by the dynamic community detection procedure; Mucha et al., 2010) and of the number of communities with extended practice, computed for all the frequency bands. *Q* is a measure of the quality of a partition into functional communities and quantifies the strength of functional modularization and the separation between functional modules. *Q* increased with increasing number of trials practiced in the theta and alpha bands (Figures S2a-b) suggesting that community structure in functional networks become more pronounced with learning. Indeed, increasing *Q* indicates that with learning the pattern of functional connectivity in the brain can be better clustered into distinct communities of brain regions exhibiting similar time courses. In the beta and gamma bands (Figures S2c-d) multilayer modularity increased or remained stable in the first ~100 trials of practice and then decreased with extended learning, suggesting a less pronounced community structure with learning in these bands. The number of communities increased with practice in all frequency bands (Figures S2e-h), apart from an initial decrease during the first ~100 trials in the beta and gamma bands (Figures S2g-h). Like for flexibility, an increasing number of communities with learning is consistent with an increased specificity of functional connectivity patterns.

| **Training Intensity** | **EEG Session 1** | **Home** Weeks 1 & 2 | **EEG Session 2** | **Home** Weeks 3 & 4 | **EEG Session 3** | **Home** Weeks 5 & 6 | **EEG Session 4** |
|---|---|---|---|---|---|---|---|

| | | | | | | | |
|---|---|---|---|---|---|---|---|
| EXT 1&2 | 50 **(50)** | 640 (690) | 50 **(740)** | 640 (1380) | 50 **(1430)** | 640 (2070) | 50 **(2120)** |
| MOD 1&2 | 50 **(50)** | 100 (150) | 50 **(200)** | 100 (300) | 50 **(350)** | 100 (450) | 50 **(500)** |
| MIN 1&2 | 50 **(50)** | 10 (60) | 50 **(110)** | 10 (120) | 50 **(170)** | 10 (180) | 50 **(230)** |

**Table S2: Number of trials completed at the end of each EEG session and every 2-weeks of home practice, displayed for each sequence practiced during the task. Cumulative values are displayed in brackets. Bold values in parentheses correspond to the EEG sessions and correspond to the values on the x-axis in Figures 2 and 4 in the main text. At the end of the fourth EEG session, participants have practiced 2120, 500 and 230 trials for the EXT, MOD, and MIN sequences, respectively**.

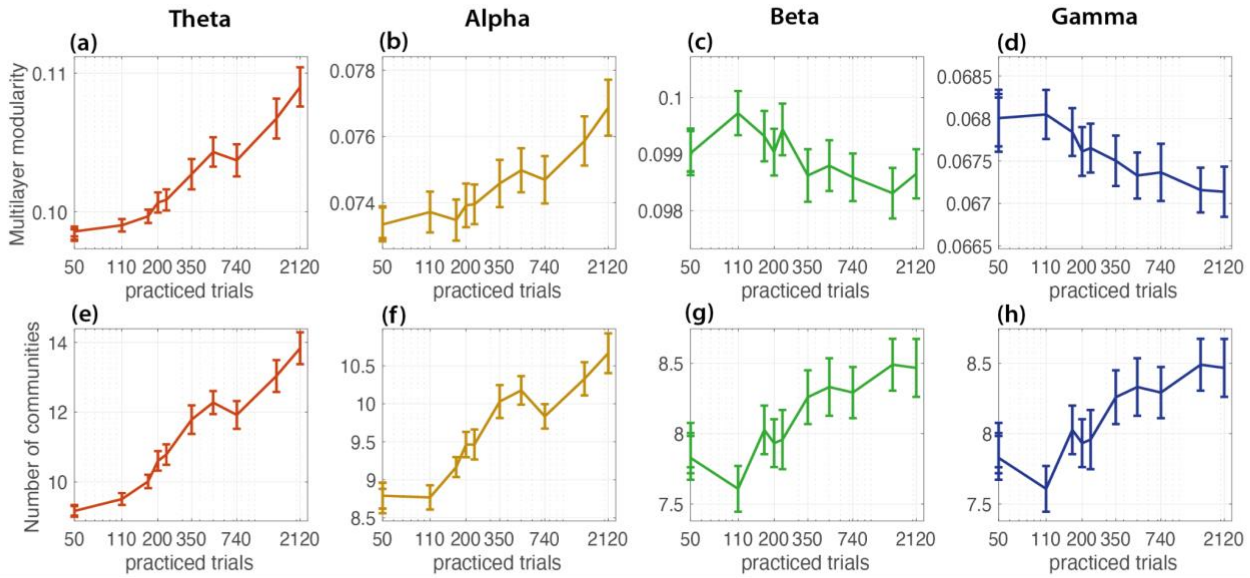


**Figure S2: Multilayer modularity and number of communities for the theta, alpha, beta and gamma band, computed from EEG data recorded during EEG sessions and displayed as a function of practiced trials (see Table S2 for the relationship between practiced trials and the corresponding trained sequence and EEG session).**

### Supplementary section 3: Functional correlates of performance

| **Control Variable** | **Lateral Orbitofrontal L** | **Parsorbitalis R** | **Rostral Anterior Cingulate R** | **Rostral Middle Frontal L** |
|---|---|---|---|---|
| ***None*** | rho = -0.50<br>p-value = 0.005 | rho = -0.37<br>p-value = 0.044 | rho = -0.44<br>p-value = 0.014 | rho = -0.50<br>p-value = 0.005 |
| ***Initial MT*** | rho = -0.44<br>p-value = 0.017 | rho = -0.34<br>p-value = 0.067 | rho = -0.37<br>p-value = 0.049 | rho = -0.40<br>p-value = 0.033 |

| | | | | |
|---|---|---|---|---|
| ***Final MT*** | rho = -0.52<br>p-value = 0.004 | rho = -0.38<br>p-value = 0.043 | rho = -0.44<br>p-value = 0.017 | rho = -0.50<br>p-value = 0.006 |
| ***Initial & Final MT*** | rho = -0.52<br>p-value = 0.005 | rho = -0.31<br>p-value = 0.109 | rho = -0.38<br>p-value = 0.047 | rho = -0.40<br>p-value = 0.036 |

**Table S3: Relationship between learning rate *K* and flexibility estimates in the beta band computed from the first EEG session. Spearman's rank correlation results are shown, including additional control for initial and final MT.**

#### Supplementary section 4: Exploration of potential confounders

Here, we shed light on possible confounding factors that could somehow influence brain network dynamics. We first investigated the power spectral density (PSD) of the EEG signals and the possibility that our flexibility assessment was in part related to a change in signal power and less to a change in the brain network dynamics. This was done although the PLV is sensitive to the phase more than the amplitude of the signal. We then studied the relationships between network dynamics and the implicit structure of the task (e.g., trial duration), and produced new estimates of flexibility by keeping the length of the analyzed signal constant. It is important to specify that this latter approach is of exploratory nature and contains several limitations. Among these, the most critical is the loss of a significant amount of EEG signal due to the proposed threshold procedure to resize the trials to the same length. More details will be provided in the following text.

##### *4.1 Relationship between EEG signal power and flexibility estimates*

We studied PSD changes with learning as we did with networks. The absolute power of the recorded EEG signals was computed for each experimental session. First, we computed a power spectral density for the signals recorded at each electrode and each trial via Welch's method (0.5 s window size and 50% overlap). We then computed the absolute power at each frequency band (theta, alpha, beta and gamma) by summing the PSD at the corresponding frequencies. We finally averaged power estimates across electrodes, and then across trials of the same intensity (EXT, MOD, and MIN). The results are displayed in Figure S3. Repeated measure ANOVAs with training intensity (EXT, MOD and MIN) and EEG session (session 1, session 2, session 3 and session 4) as within-subject factors were applied to the power estimates. A main effect of the session was observed in the beta ($F$(1.71, 49.60) = 6.07, $p$ = .006, $\eta^2_p$= 0.173) and gamma ($F$(1.53, 44.26) = 3.85, $p$ = .039, $\eta^2_p$= 0.117) bands. Post-hoc analysis revealed that power significantly decreased between session 1 and session 2 in both frequency bands (beta: Δ(S2-S1) = -1.26, SE = 0.40, t = -3.18, p = .021 Bonferroni-corrected ; gamma: beta: Δ(S2-S1) = -0.74, SE = 0.28, t = -2.65, p = .013 non-corrected). It was also observed a main effect of training intensity in the alpha (F(1.48, 42.83) = 8.55, p = .002, η2p= 0.228) and beta (F(1.53, 22.25) = 4.26, p = .029 , η2p= 0.128) bands. Post-hoc analysis revealed that spectral power was significantly higher during execution of EXT as compared to MOD and MIN sequences in the alpha band (Δ(EXT-MOD) = 0.16, SE = 0.04, t = 3.84, p = .002 Bonferroni-corrected ; Δ(EXT-MIN) = 0.24, SE = 0.07, t = 3.28, p = .008 bonferroni-corrected), while spectral power during execution of EXT sequences was significantly higher MIN sequences in the beta band (Δ(EXT-MIN) = 0.13, SE = 0.05, t = 2.42, p = .022 not-corrected). No interaction between session and training intensity was observed in any of the frequency bands.

In summary, the results presented in Figure S3 showed a significant reduction in absolute beta and gamma band power between the first and the second session followed by non-significant variations until the fourth session. Using Spearman's correlations, we identified a weak significant negative correlation with flexibility only in the beta ($r$ = -0.28, $p$ < .001) and gamma ($r$ = -0.26, $p$ < .001) frequency bands. These results indicate that the observed flexibility increase could be considered independent of signal power modulation with learning in the theta and alpha bands, and largely unrelated to signal power in the beta and gamma bands.

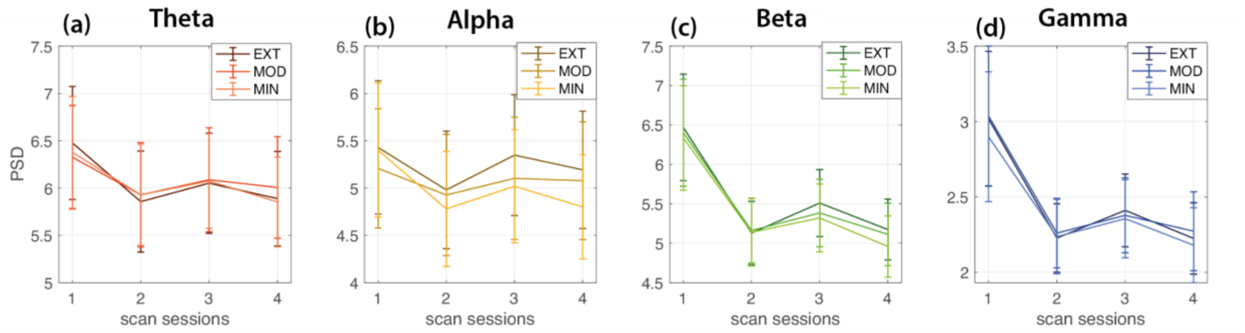


**Figure S3: (a-d) Absolute power grouped by training intensity (EXT, MOD, and MIN), computed in the (a) theta, (b) alpha, (c) beta and (d) gamma bands from EEG data recorded at consecutive EEG sessions. Error bars indicate the standard error of the mean across participants.**

***4.2 Accounting for signal length in global and regional flexibility computation***

The impact of variable signal's length on brain network estimates was recently investigated with the objective of revealing organizational principles of brain functional connectivity (Telesford et al., 2016). The authors of this work showed that changes in the length of the window used to estimate connectivity layers were related to changes in the magnitude and range of variability of estimated network statistics such as flexibility. Previous studies (Bassett et al., 2015, 2013) using the same experimental protocol of our study and performing analysis similar to those implemented in this work, did not include any supplementary analysis, leaving open the possibility that part of the results obtained could be explained by a variable length of the signal used for the construction of multilayer networks across experimental sessions. Indeed, neuroimaging studies often compute functional connections by means of pairwise wavelet coherence (as in the case of fMRI studies) or by measuring the degree of synchronization between the instantaneous phases (as in the case of EEG studies) estimated from the regional activities of all pairs of brain regions. Both approaches are not independent of the length of the signal used to compute functional connectivity estimates. In both cases, consecutive temporal layers made of connectivity matrices obtained from shorter signals could show higher across-layers variability as compared to those eventually computed from longer signals. In the context of community-based approaches, when multilayer networks are built from functional connectivity matrices computed from signal extracted at consecutive time windows, this could influence the optimization process of community assignment and the measures that depend on these dynamics, including flexibility.

We initially assessed the relationship between the duration of the trials used to compute the connectivity layers of the multilayer networks and brain network flexibility. To this end, we have extracted from our data a series of additional variables that were correlated with flexibility: (i) the average window length, expressed as the number of dynamic PLV used to compute static plv matrices; (ii) the average of the internal variance of the static PLV; and (iii) the coefficient of variation of the static PLV mean computed across layers of the multilayer temporal networks. These variables were computed for each participant, EEG session, training intensity (EXT, MOD and MIN) and frequency band (see Figures S4a-f for a representative visualization in the beta band). Spearman correlations between these variables and flexibility were computed by considering data from all participants, EEG sessions and training intensities. As there is great consistency between the results of the analysis made in each frequency band, we report correlation values for the beta band as representative for the others (Figures S5a-c). Flexibility varied as a constant power of window length, with flexibility increasing with decreasing window length ($r$ = -0.89, $p < .001$; Figure S5a). Flexibility positively correlated with the average variance of static PLV ($r$ = 0.80, $p < .001$; Figure S5b) and with the coefficient of variation of the static PLV mean computed across network layers ($r$ = 0.59, $p < .001$; Figure S5c). These results highlight that flexibility was correlated with variables related to the implicit structure of the task. Although this latter evidence does not reveal causality between the correlated variables, the results suggest that properties of multilayer networks and flexibility may have a common origin due to reduced signal length with learning.

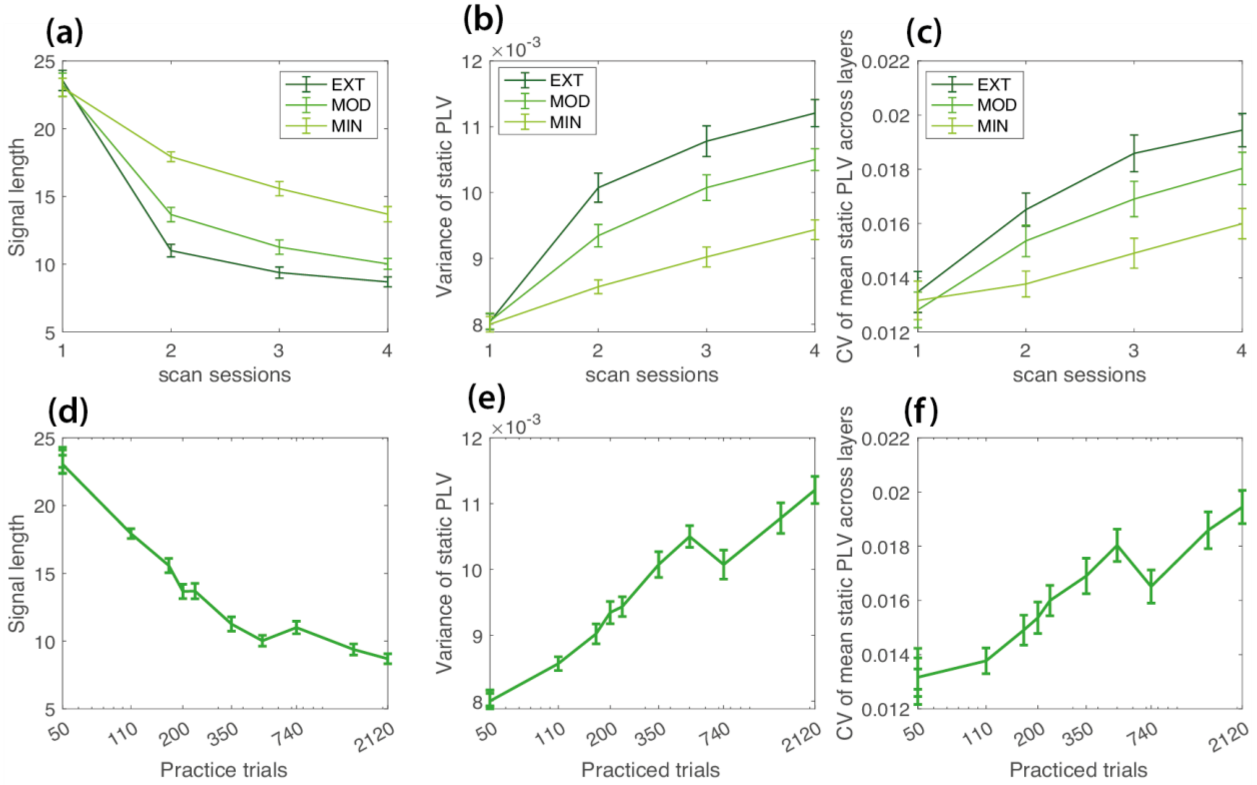


**Figure S4: (a-c) From left to right: signal length, variance of static PLV and CV of mean static PLV across multilayer networks, grouped by training intensity (EXT, MOD, and MIN) and displayed for consecutive EEG sessions. (d-f) From left to right: signal length, variance of static PLV and CV of mean static PLV across multilayer networks, computed as a function of the number of trials**

**completed after a given EEG session. Error bars indicate the standard error of the mean computed across participants. Plots are displayed for the beta band.**

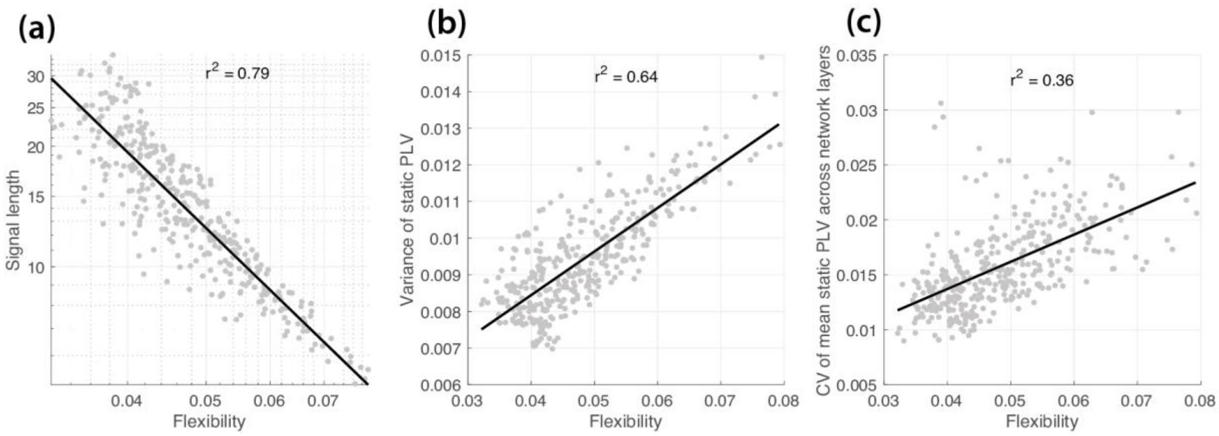


**Figure S5: Linear regression of flexibility scores with (a) the average window length, expressed as the number of dynamic plv used to compute static plv matrices; (b) the average of the internal variance of the static plv; and (c) the coefficient of variation of the static plv mean computed across layers of the multilayer temporal networks. Linear regressions were computed by considering data from all participants, EEG sessions and training intensities in the beta band.**

Finally, we estimated flexibility by keeping the length of the analyzed signal constant. We opted for a thresholding procedure of task trials that was optimized for each participant and training intensity (EXT, MOD, MIN sequences). We first defined, for each participant and training intensity, the duration of the trial performed most rapidly across the 4 EEG sessions. This duration implicitly defines the number of dynamic PLV matrices to be used for the calculation of static matrices in each of the remaining trials across the 4 sessions. The multilayer temporal networks for each training intensity were then constructed, for each EEG session, from corresponding static connectivity matrices of the same intensity. As in the previous analysis, multilayer temporal networks were analyzed through multilayer modularity optimization and flexibility used as network diagnostics. To allow comparison of flexibility estimates across participants, EEG sessions and training intensity, flexibility estimates for each training intensity were normalized with respect to the corresponding flexibility estimates obtained from the first EEG session. The above procedure was repeated for each participant and each frequency band.

##### *4.2.1 Quantification of global flexibility*

We evaluated changes in global flexibility over the six weeks of motor skills practice, for each frequency band (Figure S6). Repeated measure ANOVAs with training intensity (EXT, MOD and MIN) and EEG session (session 1, session 2, session 3 and session 4) as within-subject factors were applied to the global flexibility estimates for each frequency band (Figures S6a-d). A main effect of session was found in the theta ($F(1.97, 57.25) = 6.65$, $p = .003$, $\eta 2p= 0.187$), beta ($F(2.59, 75.01) = 2.99$, $p = 0.043$ , $\eta 2p= 0.093$) and gamma ($F(1.33, 38.63) = 3.97$, $p = 0.042$ , $\eta 2p= 0.120$) bands. In the theta band, Bonferroni-corrected post-hoc comparisons revealed that global flexibility significantly increased between the first and the second session ($\Delta(S2\text{-}S1) = 0.054$, SE = 0.017, t = 4.688, $p = .034$), followed by non-significant variations across successive sessions. In the beta and gamma bands, post-hoc comparisons revealed that global flexibility significantly increased between the first and the second session (beta: $\Delta(S2\text{-}S1) = 0.052$, SE = 0.022, t = 2.363, $p = .025$; gamma: $\Delta(S2\text{-}S1) = 0.082$, SE = 0.040, t = 2.044, $p = .05$), followed by not-significant variations across successive sessions. A main effect of training was only observed in the gamma band ($F(1.48, 42.81) = 7.91$, $p = .003$, $\eta 2p= 0.214$). Bonferroni corrected post-hoc comparisons revealed that global flexibility was significantly higher during practice of MIN sequences compared to MOD and EXT sequences ($\Delta(MIN\text{-}EXT) = 0.042$, SE = 0.012, t = 3.440, $p = .005$; ($\Delta(MIN\text{-}MOD) = 0.037$, SE = 0.014, t = 2.660, $p = .037$). A significant interaction between training intensity and session was observed in the gamma band ($F(2.99, 86.64) = 5.69$, $p = .001$, $\eta 2p= 0.164$). Post-hoc tests revealed that global flexibility increased more rapidly across scan sessions for sequences that were minimally practiced during home-based sessions as compared to moderate and extensively practiced ones ($\Delta(S2\text{-}S1)EXT = 0.063$, SE = 0.038, t = 1.671, $p = .105$; $\Delta(S2\text{-}S1)MOD = 0.073$, SE = 0.036, t = 2.051, $p = .049$; $\Delta(S2\text{-}S1)MIN = 0.109$, SE = 0.049, t = 2.254, $p = .032$). Global flexibility was not significantly modulated in the alpha band.

In Figures S6e-h the same results displayed in Figures S6a-d are shown sorted according to the number of trials performed (refer to Table S1 to see how cumulative practice trials were computed). Global flexibility seems to increase already after very few practiced trials, but the rate of growth does not seem so marked and prolonged throughout the training period as previously observed. To better quantify these variations and reduce the multiple comparisons, we averaged the global flexibility over estimates obtained at consecutive practice trials (during the first session where each sequence was practiced for 50 trials, and then across three estimates obtained between 110 and 200 trials, between 230 and 500 trials, and between 740 and 2120 trials). We then ran an additional repeated-measures ANOVA analysis on global flexibility with practice trials (first session, 110 to 200, 230 to 500, and 740 to 2120 practiced trials) as within-subject factors. A main effect of practice was found in the theta ($F(1.41, 87) = 5.07$, $p = .019$, $\eta^2_p = 0.076$), beta ($F(1.80$ $52.08) = 4.89$, $p = .014$, $\eta^2_p = 0.144$), and gamma (F(1.16, 33.72) = 4.66, p = .033 , η2p= 0.139) bands. In the theta band, Bonferroni corrected post-hoc comparisons revealed that global flexibility around 200 to 500 practiced trials was significantly higher than pre-practice trials estimated during the first EEG session (Δ(230-500 vs first session) = 0.058, SE = 0.019, t = 3.050, p = .029) without significant variations with increasing practice. In the beta band, Bonferroni corrected post-hoc comparisons revealed that global flexibility around 100 to 200 practiced trials was significantly higher than pre-practice trials estimated during the first EEG session (Δ(110-200 vs first session) = 0.065, SE = 0.021, t = 3.094, p = .026) without significant variations with increasing practice. In the gamma band, Bonferroni corrected post-hoc comparisons revealed that global flexibility around 100 to 200 practiced trials was significantly higher than pre-practice trials estimated during the first EEG session (Δ(110-200 vs first session) = 0.101, SE = 0.023, t = 3.023, p = .026), but then significantly decreased with increasing practice (Δ(740-2120 vs 230-500) = -0.031, SE = 0.010, t = -2.974, p = .035). Global flexibility was not significantly modulated in the alpha band.

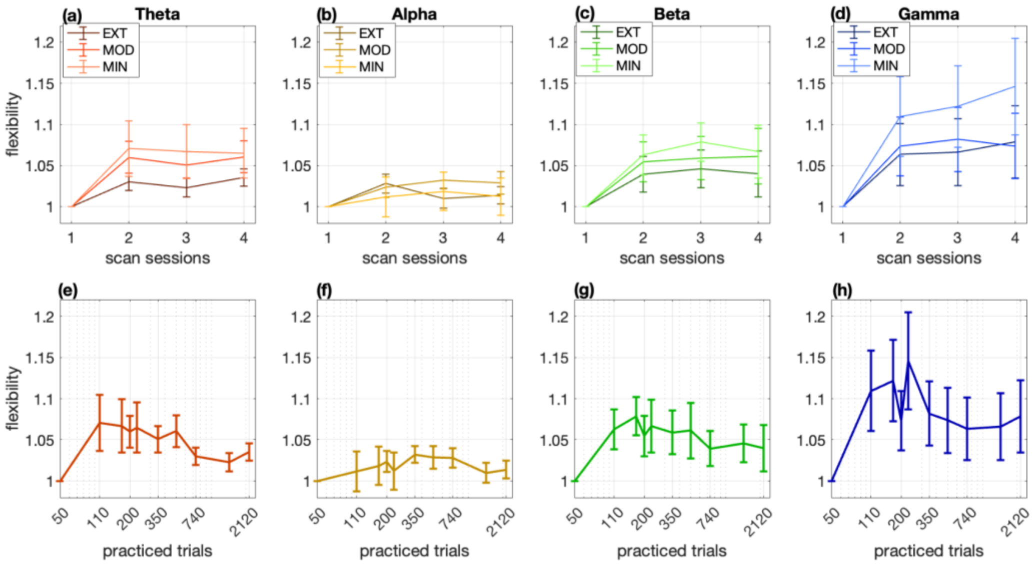


**Figure S6: Global flexibility changes with learning after accounting for signal length. (a-d) From left to right: global flexibility, grouped by training intensity (EXT, MOD, and MIN), computed in the**

**theta, alpha, beta and gamma bands from EEG data recorded at consecutive EEG sessions. (e-h) From left to right: global flexibility computed as a function of the number of trials completed after a given EEG session. Error bars indicate the standard error of the mean across participants. Flexibility estimates for each training intensity and each participant were computed as relative variations with respect to the corresponding estimates computed from data of the first EEG session.**

##### *4.2.2 Quantification of regional flexibility*

Most of the significant differences in overall flexibility, obtained after correcting for the decrease of signal length with practice, were expressed in a marked manner already after only a few practiced trials (mostly between 110 and 200 trials) in three of the observed frequency bands. This aspect represents a key difference with respect to the previous analysis (Figure 4) where global flexibility was shown to constantly increase with practice. In Figure S7, we highlight the regions that most consistently show an increase of flexibility with practice. Contrasts are shown between 110 and 200 practice trials and the first scan in the theta (Figure S7a), beta (Figure S7c), and gamma (Figure S7d) bands. Although global flexibility did not vary significantly with overall practice in the alpha band, we report in Figure S7b the regional contrasts between flexibility estimated between 230 and 500 trials and the first scan. This training level was chosen because the flexibility difference with respect to the first session $\Delta(230-500 \text{ vs first session})$ was maximal. Similar to Figure 5, in Figure S8 we display regions with the most significant increase in flexibility between the first and the last EEG session.

In the theta band (Figure S8a), similar to the results shown in Figure 5a, flexibility increased in regions of the prefrontal (pars opercularis and triangularis, right medial orbito-frontal, left pars orbitalis, and left rostral middle- and lateral orbito-frontal), limbic (right caudal anterior cingulate, and left anterior cingulate), centro-parietal (supra marginal, precuneus, left postcentral, right precentral, paracentral and inferior parietal), and temporal (inferior and right superior) lobes. Fewer regions displayed significant contrasts in the alpha, beta and gamma bands. In the alpha band (Figure S8b), flexibility increase was observed in centro-parietal (precentral and paracentral, right superior and inferior parietal) regions and in the posterior cingulate, in partial overlap with the results shown in Figure 5b. Significant increase was also observed over central regions (pre and postcentral) in the beta and gamma bands (Figure S8c and S8d, respectively). Flexibility increases in the right superior and left middle temporal regions were specific to the beta band.

Concerning the beta and gamma bands, we did not observe a clear regional overlap with the results obtained without correction (Figures 5c and 5d).

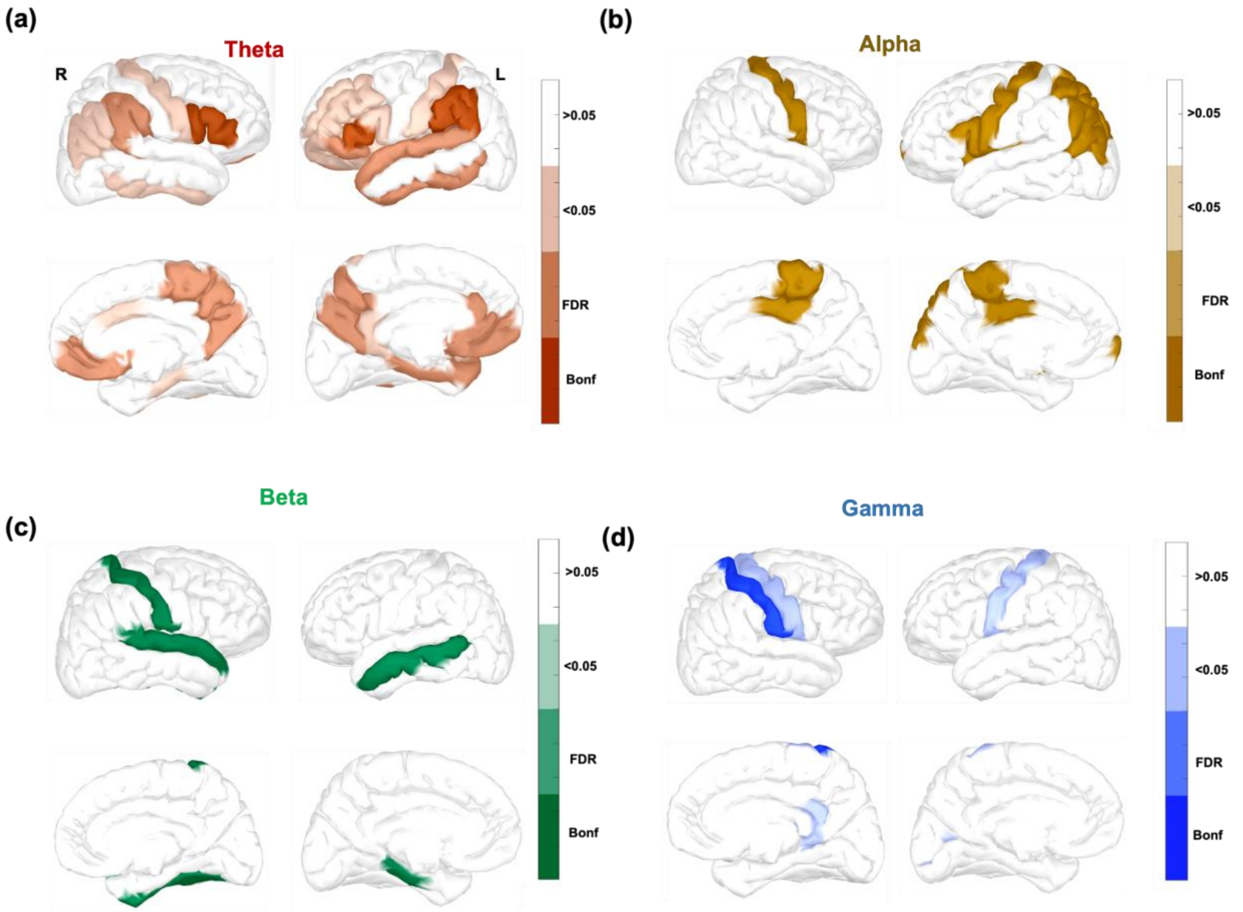


**Figure S7: Brain regions displaying the most significant increase in flexibility between EEG recordings in session 1 and recordings occurring between 110 and 200 practiced trials for the (a) theta, (c) beta, and (d) gamma frequency bands, and between 230 and 500 trials for the alpha (b) band. Contrasts are assessed using a Wilcoxon test. Brain regions displaying a significant increase in flexibility after Bonferroni and FDR correction are displayed in dark and medium dark colors. Light colors are used for brain regions with *p* < 0.05 that did not survive the Bonferroni or FDR correction. Brain regions with non-significant changes are displayed in white.**

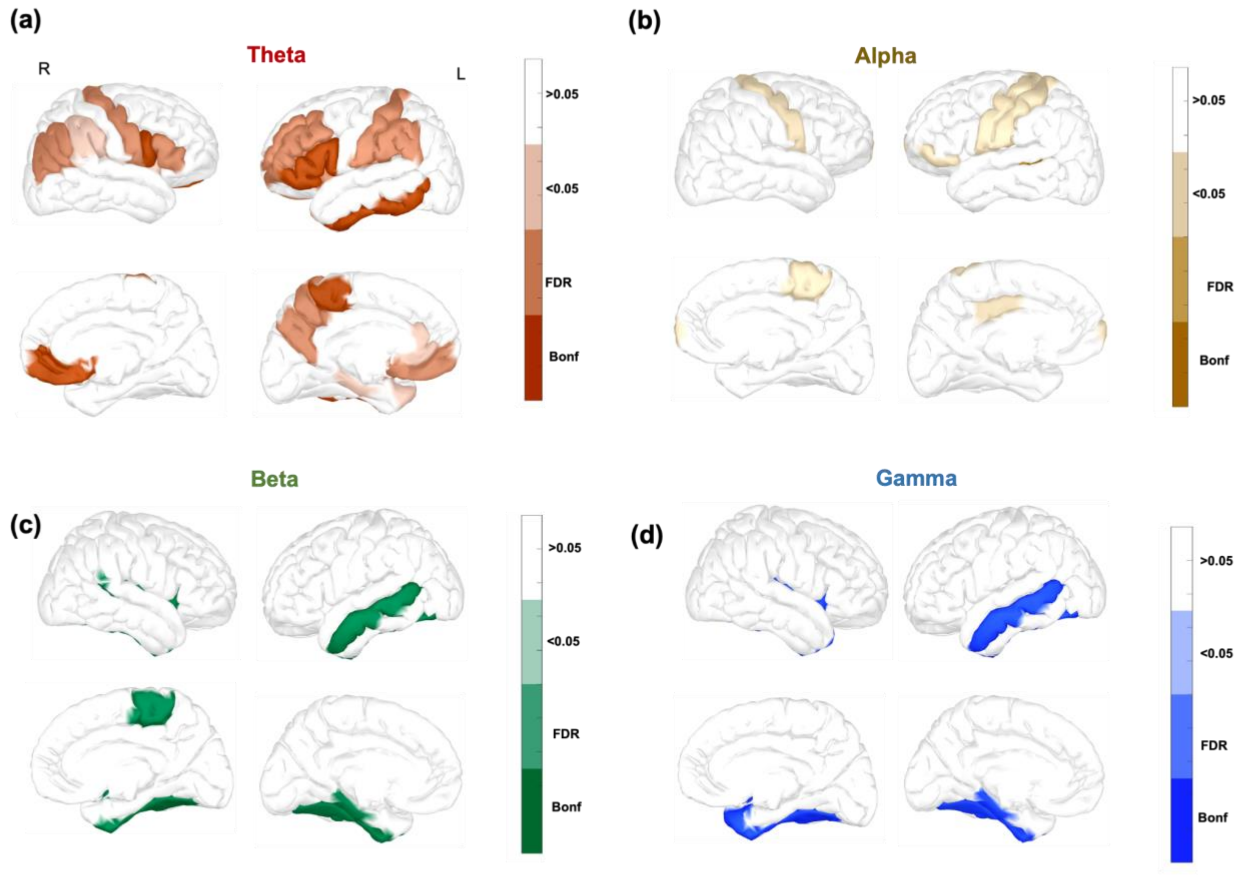


**Figure S8: Brain regions displaying the most significant increase in flexibility between EEG recordings in session 1 and 4 for the (a) theta, (b) alpha, (c) beta, and (d) gamma frequency bands. Contrasts are assessed using a Wilcoxon test. Brain regions displaying a significant increase in flexibility after Bonferroni and FDR correction are displayed in dark and medium dark colors. Light colors are used for brain regions with $p < 0.05$ that did not survive the Bonferroni or FDR correction. Brain regions with non-significant changes are displayed in white.**
